# *Nicastrin* haploinsufficiency alters expression of type-I interferon-stimulated genes in two immortalized human cell lines

**DOI:** 10.1101/353482

**Authors:** Li Cao, David J. Morales-Heil, Elisha D. O. Roberson

**Author notes:** Corresponding author: Elisha D. O. Roberson, Ph.D. Washington University Depts. of Medicine and Genetics Division of Rheumatology 660 South Euclid Ave. Campus Box 8045 St. Louis, MO 63110 United States.

## Abstract

A.

**Background:** Hidradenitis suppurativa (**HS**) is a chronic skin disease. The symptoms can be severe, and include intensely painful nodules and abscesses in apocrine-gland rich inverse skin, such as the buttocks, under the arms, and the groin. Autosomal dominant forms of HS exist, but are rare. Some of these kindred have heterozygous loss-of-function rare variants in the γ-secretase complex component *nicastrin (NCSTN).*

**Objectives:** We wanted to know what effect *NCSTN* haploinsufficiency has on human keratinocytes to assess potential mechanisms for lesion development.

**Methods:** We knocked down *nicastrin* using an shRNA construct in both a keratinocyte cell line (HEK001) and an embryonic kidney cell line (HEK293). We assessed differential gene expression using RNA microarray. We also generated a *NCSTN* heterozygous deletion in the HEK293 line using CRISPR/Cas9 genome-editing and assessed NFKB activity in this line using a luciferase reporter.

**Results:** The keratinocyte *NCSTN* knockdown cell line demonstrated significantly increased expression of genes related to the type-I interferon response pathway when compared to controls. Both HEK001 and HEK293 knockdowns demonstrated evidence for altered growth. We observed a small, but significant increase in NFKB signaling in response to TNF treatment a HEK293 line genome-edited for reduced *NCSTN*.

**Conclusions:** Our data suggest a role for increased keratinocyte inflammatory responsiveness in familial HS. Confirming this phenotype, and characterizing additional effects in different cell types, will require study beyond cell lines in primary cells and tissues.

## B. Introduction

Hidradenitis suppurativa (**HS**), also known as acne inversa, is a severe and disfiguring skin disease. The prevalence in Caucasians is estimated to be as high as 1%, making it as prevalent as well-recognized skin diseases such as psoriasis and atopic dermatitis^1,2^. The first onset of symptoms typically occurs after puberty, in the teens and early 20s. The disease course is chronic with periodic relapses. The most frequently affected areas are the axillae, groin, pubic area, and inner thighs^3^. Lesions can affect the perianal region, as well as the submammary and inframammary folds. The mildest form of HS consists of inflamed and painful nodules or lumps. In more severe forms of HS, the painful lesions progress to full abscesses. Multiple, adjacent abscesses form communicating subcutaneous sinus tracts that drain a purulent, malodorous fluid. The chronic course and severity of lesions leads to hypertrophic scarring. These lesions are intensely painful and significantly reduce quality of life in affected individuals^4–7^.

In 1985, Fitzsimmons and Guilbert showed evidence for a Mendelian-like pattern of autosomal dominant HS in some families^8^. Between 20-40% of individuals with sporadic HS ascertained from multiple populations report having at least one affected relative as well, suggesting an appreciable genetic risk^4,9–11^. One of the first strong linkage regions was mapped to chromosome 1 between *D1S248* and *D1S2711* in three generations of a Chinese kindred with autosomal dominant HS (hg19 g.chr1:107154920-185518094; 78 Mb)^12^. Rare genetic variants in components of the γ-secretase complex have since been discovered in some kindred with familial HS (**fHS**). Active γ-secretase requires nicastrin (NCSTN), anterior pharynx-defective 1 (APH1A or APH1B), presenilin enhancer gamma secretase subunit (PEN2), and a presenilin (PSEN1 or PSEN2). Both *NCSTN* and *APH1A* fall within the initially reported linkage region. Variants linked to fHS have been identified in *PSEN1, PSENEN* (the PEN2 gene), and *NCSTN* (OMIMs #142690, #613736, #613737)^13–22^. The majority of these variants are in *NCSTN*, and the same premature termination variant has even been reported in both a Chinese family and an African American family with autosomal dominant HS^13,14^. Novel variants have also been discovered in sporadic cases with no self-reported family history of disease, but the significance of these variants is not yet clear^9^. In some studies only a minority of fHS kindred (20-30%) have an identified loss-of-function variant in γ-secretase components^17,18^. This suggests that much of the genetic contribution to fHS has therefore not been identified, further confounding both the identification of loss-of-function variants in sporadic HS cases and the development of therapeutics.

The γ-secretase complex cleaves type-1 single-pass transmembrane proteins. Stable complex formation requires a presenilin to be stabilized by APH1 and NCSTN, followed by interaction with PEN2^23,24^. Yeast, which lack γ-secretase, can form active γ-secretase complexes with exogenous overexpression of the individual components^23^. Substrates for the complex have both an intracellular (cytoplasmic domain) and extracellular (ectodomain) portion. γ-secretase can only act on proteins with a shortened ectodomain, typically due to cleavage by a metalloproteinase. γ-secretase can then cleave in the substrate’s transmembrane domain, releasing the cytoplasmic domain within the cell and the ectodomain into the extracellular space. The cleaved cytoplasmic domain may then translocate to the nucleus if it contains appropriate nuclear localization signals and activate gene expression. Due to the high number of potential substrates with diverse molecular functions, predicting the effect of *NCSTN* haploinsufficiency is difficult.

The *NCSTN* rare variants so far identified in fHS suggest a haploinsufficiency phenotype. Early stops and frameshift variants would likely lead to non-sense mediated decay, supported by lower *NCSTN* expression in peripheral blood from individuals carrying such rare variants^13,17^. To assess the biological impact of *NCSTN* haploinsufficiency we used shRNA to knockdown *NCSTN* in a human epidermal keratinocyte cell line (**HEK001**) and a human embryonic kidney line (**HEK293**), and profiled the transcriptomes via microarray. We later generated a heterozygous *NCSTN* deletion in HEK293 cells. Our overall approach is diagrammed in **Supp. Fig. 1**. Surprisingly, we discovered that reduced nicastrin caused disrupted interferon signaling in both cell lines after transduction with lentivirus and selection for knockdown cells, which raises the question of what systemic effects exist for individuals with familial HS.

## C. Materials & Methods

### C.1. Cell culture

We obtained HEK001, HEK293, and HEK293T lines from the ATCC, and cultured them at 37°C with 5% CO_2_. We cultured the HEK293 and HEK293T cells in Dulbecco’s Modified Eagle Medium (**DMEM**) supplemented with 10% fetal bovine serum, 2 mM glutamine, 100 units/mL penicillin and 100 μg/mL streptomycin. We cultured HEK001 cells in Keratinocyte-SFM (serum-free medium) supplemented with human recombinant Epidermal Growth Factor and Bovine Pituitary Extract (Life Technology #17005042).

### C. 2. shRNA transfection

We transfected HEK293T cells (40-70% confluence) using the TransIT-LT1 system according to the manufacturer’s instructions (Mirus). The transfection mix contained 1 μg of shRNA hairpin plasmid and 1 μg of an 8:1 ratio of packaging plasmid (pCMF-dR8.2) and envelope plasmid (pCMV-VSV-G). After 24 hours, we removed the transfection mix and switched to 6 mL of fresh media. After another 24 hours, we collected the media, centrifuged at 1250 xg for 5 minutes, and saved the supernatant containing lentivirus.

We infected HEK001 or HEK293 cells (~70% confluence) with the lentivirus containing supernatant diluted 1:5 with complete cell culture media plus polybrene at a final concentration of 10 μg/mL. After 24 hours, we discarded the media, and replaced it with media containing puromycin. We selected HEK293 cells using 2 μg/mL puromycin and HEK001 with 0.5 μg/mL. When cultures were nearing confluence, we split them 1:10 with fresh media containing puromycin (twice in two weeks). We extracted RNA from the final cultures using the miRNeasy mini kit (Qiagen, #217004) according to the manufacturer’s instructions.

### C. 3. RT-qPCR

We generated cDNA using Superscript II according to the suggested manufacturer guidelines. This included using the included buffers and dNTP mixes with 6 μM random hexamer primers, 200 U of reverse transcriptase (**RT**), and up to 1 μg of total RNA. We incubated samples at 25°C for 10 minutes to anneal primers, 55°C for 1 hour for synthesis, and heat inactivated the RT at 70°C for 10 minutes. For RT-qPCR, we used 0.6 ng of cDNA as template with 1X SYBR Green, and 100 nM each forward and reverse primer. The reactions were incubated for 10 minutes at 95°C for initial denaturation, had up to 40 cycles of 30 second denaturation at 95°C, 45 seconds at 65°C for annealing, and 30 seconds at 78°C for extension, followed by a final 10 minute 78°C extension, and hold forever at 4°C. For relative expression, we ran at least 3 replicates of each sample-amplicon, and used the delta-delta Ct method compared to 18S rRNA to determine relative expression. We exported the quantification cycle (**Cq**) of each well and propagated standard deviation to determine the final relative expression difference. The primer sequences and details of the error propagation procedure for relative expression changes can be found in the supplementary material.

### C. 4. Microarray differential expression analysis

We analyzed the microarray data with the Bioconductor (v2.36.2) beadarray package (v2.26.1) using R (v3.4.0) and R Studio (v1.0.143). We used non-normalized, background subtracted array data as input, keeping only probes with a detection p-value < 0.05 in at least three samples. We shifted the distribution of each array by adding or subtracting a constant such that the minimum intensity value was 2.0, since background subtraction can result in negative intensities. We transformed the shifted data by log_2_, and normalized by quantile normalization. We determined differential expression using the limma (v3.32.5) package with the specified model (*NCSTN* knockdown - luciferase knockdown). We considered a gene differentially expressed if the false-discovery rate corrected p-value was less than 0.05.

We used the gProfileR g:COSt tools to analyze for pathway enrichment^25^. We included genes with a significant corrected p-value and a minimum absolute fold-change of 1.50 in pathway analyses, and analyzed genes with decreased expression separately from those with increased expression. We entered genes by RefSeq NM number rather than the gene symbol to avoid ambiguity with gene names. We ran all enrichments using the following options: significant enrichments only, run as ordered query, no electronic GO annotations, medium filtering, 0.05 p-value cutoff, maximum size of functional category of 500, g:SCS p-value correction, statistical domain including only annotated genes, and numeric IDs treated as ENTREZ gene accession numbers. The array data is publicly available in the Gene Expression Omnibus (**GEO**; GSE57949).

### C. 5. NFKB luciferase assay

We plated the HEK293 cells at a density of 100,000 cells per well in a 24-well plate 24 hours prior to transfection. We transfected cells using the transIT-LT 1 system (described above) with NFKB-luc (firefly luciferase construct containing proximal NFKB binding sites) and renilla luciferase (pGL4 as a transfection control) at a ratio of 24/1. Four hours later we treated some cultures with TNF (Sigma #T0157) at 20 ng/mL. Twenty-four hours post-transfection, we harvested the cells and assayed both firefly luciferase and renilla luciferase activity using the Dual-Luciferase Reporter Assay System (Promega). We calculated statistical significance for NFKB induction (firefly / renilla) in R using Tukey Honest Simple Differences of the ANOVA model.

### C. 6. Growth curves

We plated cells at 3,000 cells/well in standard complete culture medium (see **Cell culture**) and measured proliferation starting the second day for up to 7 consecutive days using a colorimetric assay according to the manufacturer’s instructions (CellTiter 96 Non-Radioactive Cell Proliferation Assay). We detected the color change by absorbance at 570 nm in 96-well plate reader (uQuant). Each day’s measurement had a background measurement subtracted. We calculated within day / within cell type significance by two-sided t-test.

### C. 7. Genome editing

We generated *NCSTN* knockouts in HEK293 cells with CRISPR/Cas9 based genome-editing. We modified the guide RNA scaffold to include a T>A swap (Flip) and an extended stem (Extension), called the Flip + Extension (**F+E**) guide, as previously described^26^. We targeted *NCSTN* with guide sequence GGCATACTGTACAACATAAG (g.chr1:160326441-160326460; hg19). We ordered the modified gRNA scaffold with our guide as an IDT gBlock for blunt-end cloning with the pCR-Blunt II-TOPO vector (**Supp. Table 2**). We amplified the guide RNA stock using PCR to obtain sufficient amounts for cloning. We also ordered a donor gBlock to insert a stop-codon in *NCSTN* in case homology-directed repair was favored over non-homologous end-joining (**Supp. Table 3**). We amplified the donor using PCR to prior to transfection as well. We plated HEK293 cells at a density of 350,000 cells/well in 6-well plates the day prior to transfection. We then transfected them using the LT-1 system with 0.75 μg each of humanized Cas9 plasmid (Joung lab; AddGene JDS246, plasmid #43861), guide RNA, and a donor amplicon.

After 24 hours incubation, we plated the cells in fresh media at 50,000 cells / 15 cm culture plate to dilute cells into single-cell clones. We picked samples from clonal clusters one week later, and transferred them to individual wells in a 96-well plate. We sampled a scraping from individual wells, and amplified the target region by PCR. One sample (clone 48) had a potential frameshift by Sanger sequencing. RT-qPCR confirmed less *NCSTN* mRNA compared to wildtype HEK293 cells.

## D. Results

### D. 1. *NCSTN* mRNA can be reduced to less than 50% wildtype levels using shRNA

We tested five separate shRNA constructs (A06-A10; **Supp. Table 4**) for their ability to knockdown *NCSTN* mRNA levels in the HEK001 and HEK293 cell lines (**Fig. 1**). We used two different *NCSTN* amplicons for each shRNA and tested for relative expression differences using realtime quantitative PCR (**RT-qPCR**). Only shRNA constructs A08 and A09 demonstrated consistent, strong reduction of *NCSTN* mRNA for each amplicon in both cell lines. shRNA A08 demonstrated the strongest knockdown and was used for further study. We generated stable knockdowns in both HEK001 and HEK293 for A08 and a control shRNA (shLuc; targeting luciferase). We generated each stable line in triplicate and selected for retention of the shRNA plasmid for two weeks in puromycin. We confirmed the knockdown (**Supp. Fig. 2**) and used the shLuc and shA08 line RNAs for array-based transcriptome profiling. We later generated an shGFP (shRNA targeting green fluorescent protein) stable knockdown for use in luciferase assays.

**Fig. 1.**
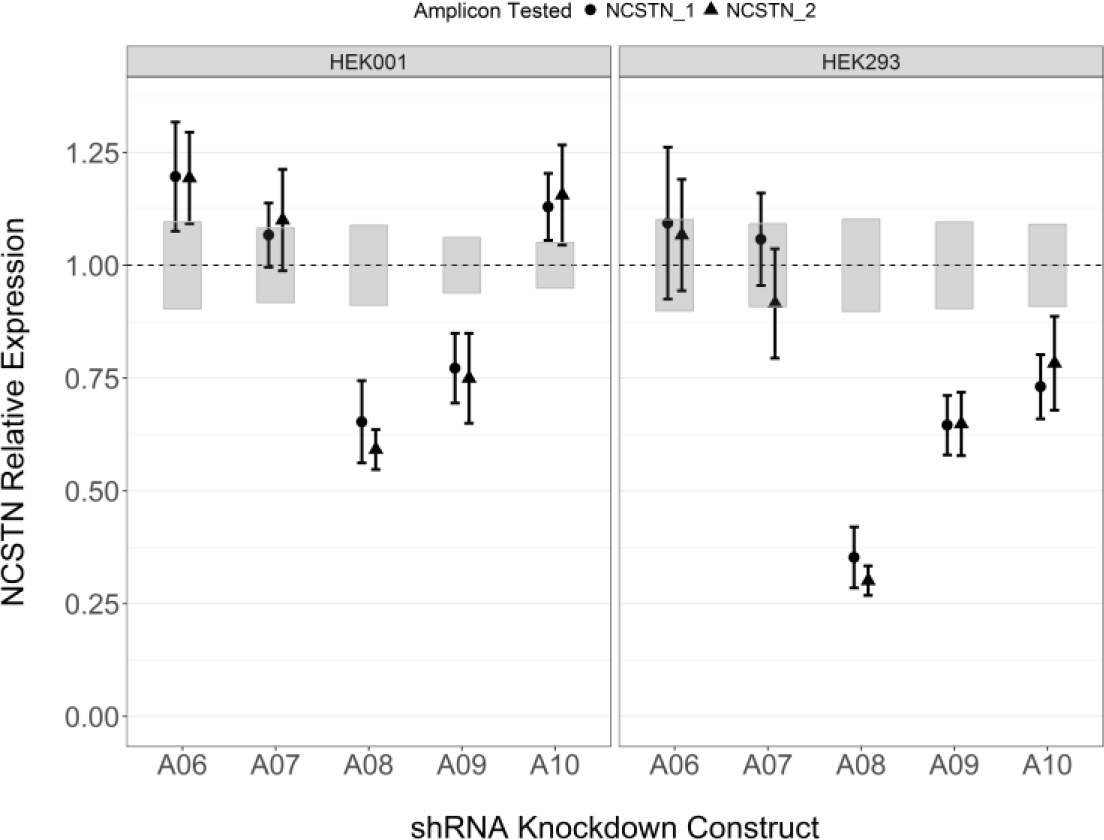
*NCSTN* shRNA construct RT-qPCR. Shown are the relative expression results for the 5 shRNA constructs (A06-A10). The x-axis is the shRNA construct used, and the y-axis is the relative expression to non-transduced controls. Points indicate mean relative expression and error bars represent standard deviation. Grey boxes indicate the standard deviation of the control cells. We tested two different *NCSTN* amplicons (circle and triangle) and each tested amplicon had 3 technical replicates. A08 had the most consistent knockdown effect.

### D. 2. HEK001 keratinocyte *NCSTN* knockdown lines have increased type-I interferon activation and disruption of cell cycle related genes

HEK001 knockdown of *NCSTN* resulted in the differential expression of 1,548 probes for 1,393 unique genes (**Fig. 2A**; **Supp. Table 8**). We confirmed a subset of these by RT-qPCR (**Supp. Fig. 3**). We focused on those genes with an adjusted p-value < 0.05 and an absolute fold-change (FC) of at least 1.50 (n=364 increased, n=324 decreased). Significant fold-changes ranged from −8.04 (nicastrin, *NCSTN*) to 5.58 (fibroblast growth factor receptor 3, *FGFR3*).

**Fig. 2.**
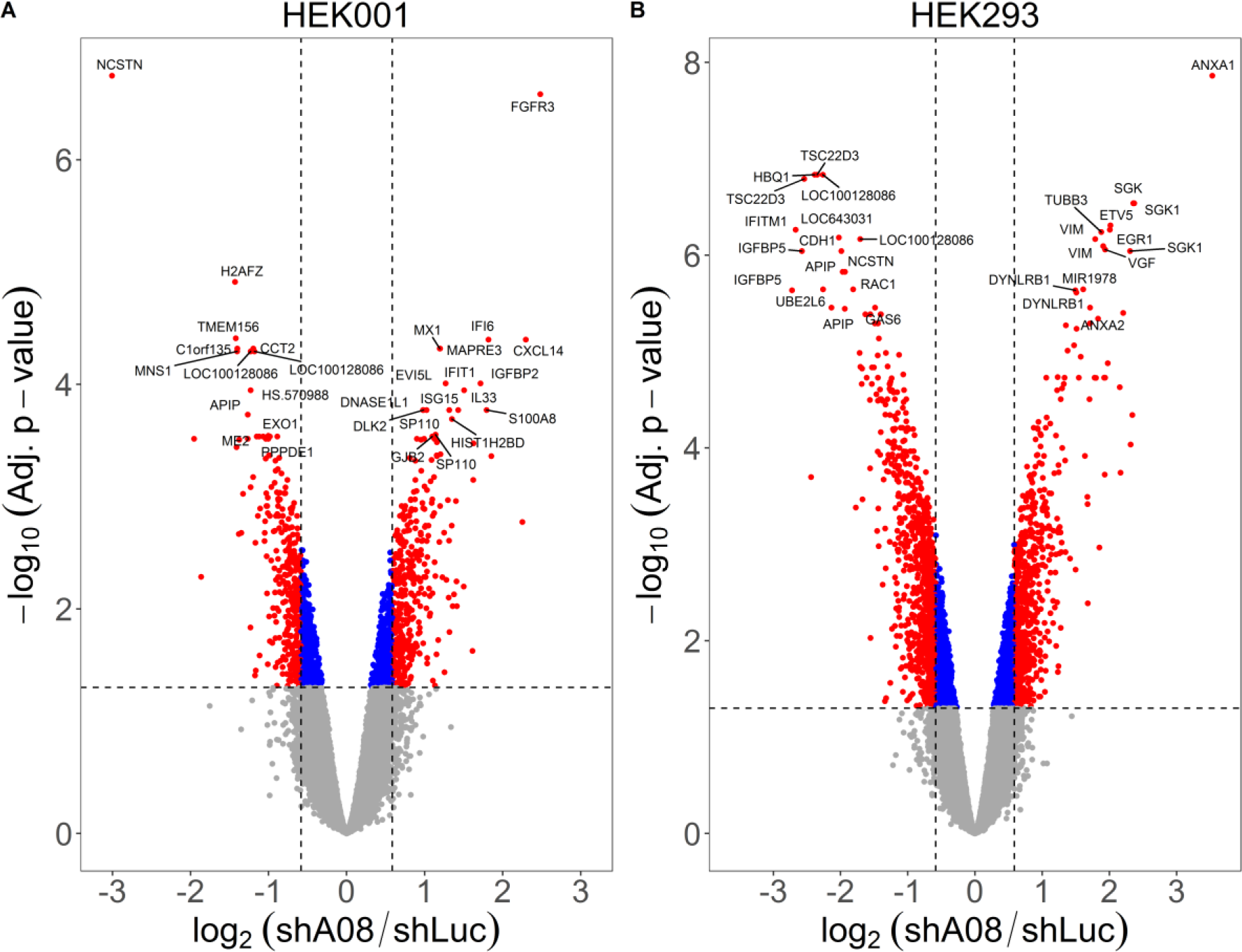
Volcano plot of differentially expressed genes with knockdown. Volcano plots for the HEK001 (A) and HEK293 (B) knockdowns. The effect size is shown on the x-axis (log2 fold-change) and the adjusted significance on the y-axis. Some top genes are individually labeled. The horizontal line represents adjusted p-value of 0.05, and the vertical lines represent 1.5-fold changes.

We tested significant genes with at least a −1.5-fold decreased expression for enrichment of known pathways. We found enrichment of genes related to cell cycle (**Supp. Fig. 4A**; **Supp. Table 9**). Decreased genes in this category included histone proteins (H2A histone family member Z, *H2AFZ;* −2.69 FC), topoisomerase 2 (DNA topoisomerase II alpha, *TOP2A;* - 1.99 FC), and centromere proteins (centromere protein V, *CENPV;* −1.98 FC), among others. We separately tested the genes with at least a 1.5-fold significantly increased expression for enrichment. Instead of cell cycle related functions, we found enrichment for type-I interferon responses (**Supp. Fig. 4B**; **Supp. Table 10**). Among the most increased genes are several associated with inflammation, including chemokines (C-X-C motif ligand 14, *CXCL14;* 4.91 FC), cytokines (interleukin 33, *IL33;* 2.69 FC) and interferon inducible proteins such as interferon alpha-inducible protein 6 (*IFI6*, 3.52 FC), and interferon-induced GTP-binding protein Mx1 (*MX1*; 2.29 FC).

### D. 3. HEK293 *NCSTN* knockdowns have decreased NOTCH and interferon signaling

The HEK293 cell lines exhibited more transcriptome changes in response to knockdown of *NCSTN.* Overall, 3,089 probes for 2,813 genes were significantly differentially expressed (**Fig. 2B**; **Supp. Table 11**). More than 1,300 genes exhibited at least an absolute 1.5-fold change (n=607 up, n=784 down), and the largest changes ranged from −6.6 FC (insulin-like growth factor binding protein 5, *IGFBP5*) to 11.6 FC (annexin a1, *ANXA1*). Again, we confirmed a subset of changes by RT-qPCR (**Supp. Fig. 5**).

As before, we tested for enrichment of known molecular pathways. Genes decreased in the HEK293s were enriched for small molecule and steroid biosynthesis (**Supp. Fig. 6A**; **Supp. Table 12**). There were other significant enrichments for NOTCH related signaling, including general NOTCH signaling, NOTCH receptor processing, and NOTCH3 signaling. The genes linked to these pathways were *NCSTN* (−3.82 FC), jagged-2 (*JAG2;* −-2.51 FC), *NOTCH3* (−3.06 FC), and *MYC* (−1.26 FC). Of note, while NOTCH related signaling was identified as an enriched pathway as a result of these transcript changes, we did not detect differences in *HEY1* (−1.06 FC; adj. P=0.83) or *HES1* (1.05 FC; adj. P=0.79), canonical immediate downstream genes regulated by NOTCH signaling. *HES5* is another of these targets, but was filtered out as it didn’t meet the detection p-value criteria. Thus, while *NCSTN* depletion resulted in decreased NOTCH pathway components, we did not see evidence for differences in downstream NOTCH signaling. In contrast to the HEK001 cells, the HEK293 genes with decreased expression were enriched for interferon signaling. This included decreases in *MX1* (−1.63 FC), *IFI6* (−2.25 FC), *IFIT1* (-3.84 FC), *IFITM1* (interferon-induced transmembrane protein 1; −6.36 FC), *IFITM2* (−3.20 FC), and *IFITM3* (−1.84 FC).

The genes with increased expression were primarily enriched for categories related to senescence, HDAC activity, and methylation (**Supp. Fig. 6B**; **Supp. Table 13**). The senescence & methylation enriched genes primarily related to histone cluster 1 and 2 genes *(HIST1H1C*, 2.37 FC; *HIST1H2AC*, 1.92 FC; *HIST1H2BD*, 2.30 FC; *HIST1H2BK*, 1.74 FC; *HIST1H4H*, 3.25 FC; *HIST2H2AA3*, 2.06 FC; *HIST2H2AA4*, 1.96 FC; *HIST2H2AC*, 1.81 FC; *HIST2H4A*, 2.50 FC; *HIST2H4B*, 2.43 FC).

### D. 4. *NCSTN* knockdown alters cell metabolism after exponential growth

The HEK001 knockdowns had decreases in cell cycle pathways and the HEK293 knockdowns had increases in senescence / cell death pathways. The convergence on cell division and senescence made us interested potential growth defects in the cells. We assayed cell proliferation in both HEK001 and HEK293 lines using a colorimetric assay. There was no statistical difference for knockdowns in either cell line during the initial exponential growth phase (**Fig. 3**). However, knockdowns of both lines demonstrated significantly reduced signal in the post-exponential phase (**Supp. Table 5**). It’s worth noting that the non-radioactive assay we used relies on conversion of an input substrate into a product with different absorbance spectra. Therefore the significantly reduced activity demonstrates reducedconversion, which could be driven by increased cell death, increased rates of senescence, or a mixture of the two.

**Fig. 3.**
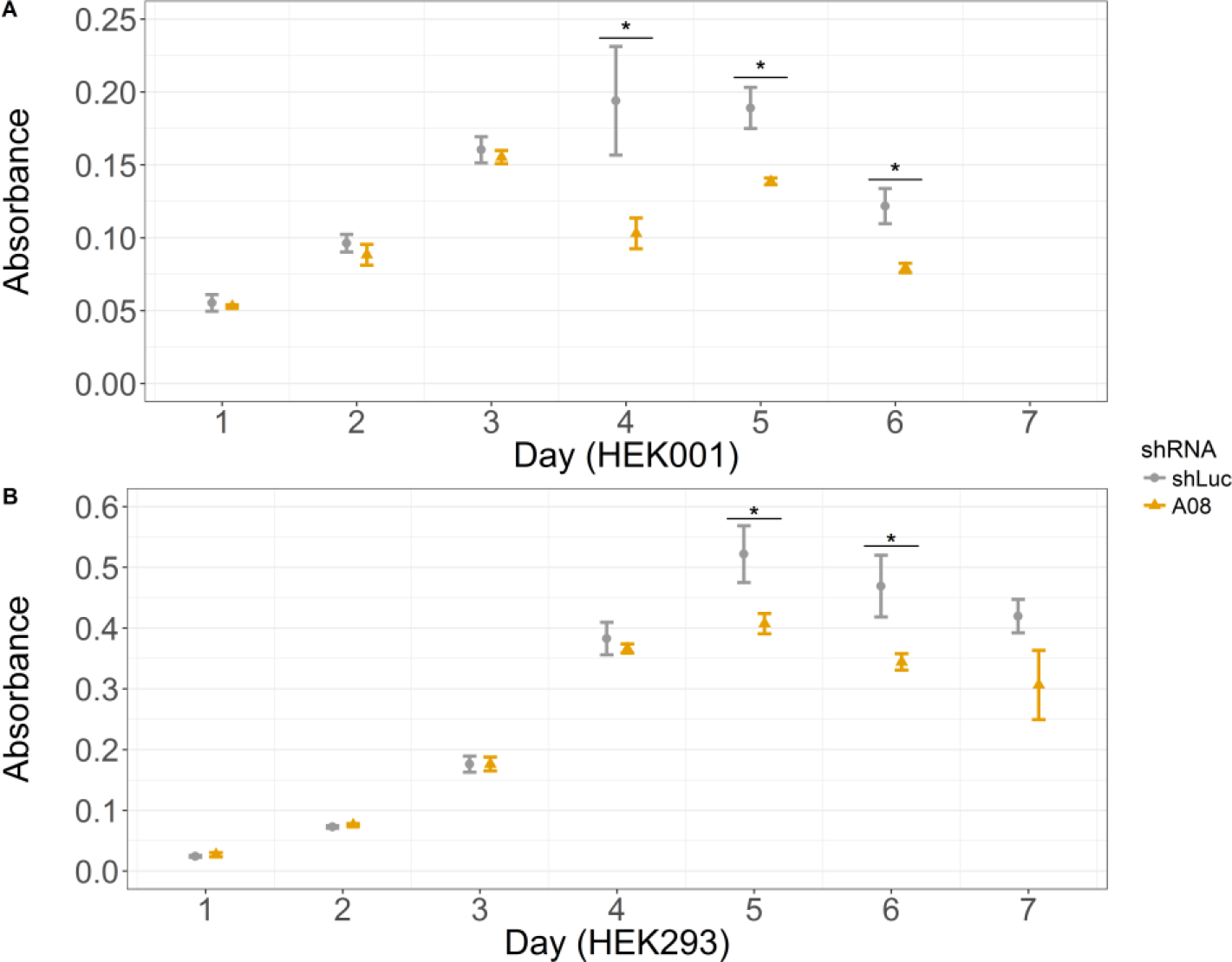
Proliferation assay reagent conversion disrupted in *NCSTN* knockdowns. We tested knockdowns in both cell types for proliferation by colorimetric substrate conversion assay. The x-axis is the day post plating and the y-axis is the background-subtracted absorbance. The shRNAs are differentiated by shape and color. The points represent the mean and error bars are the standard deviation. Each was based on 3 replicates. Any differences within a time point by t-test are indicated by solid horizontal lines with asterisks. Three late HEK001 and two late HEK293 time points are slightly but significantly lower

### D. 5. HEK293 cell lines have altered NFKB activation with decreased *NCSTN*

In sporadic HS, suppressing TNF with anti-TNF monoclonal antibodies can be an effective treatment. Given the misregulation of inflammatory genes in our *NCSTN* knockdown cell lines, we were curious whether a pathway downstream of TNF, particularly the NFKB response, was activated in our cells. To assess the effect of *NCSTN* depletion on NFKB signaling, we transfected our HEK293 knockdown cells with a plasmid containing luciferase under the control of an NFKB inducible promoter. We tested three stable knockdowns (A08, shLuc, shGFP) with no treatment and after stimulation with TNF (**Fig. 4**; **Supp. Table 6**). The A08 line demonstrated significantly increased NFKB activity compared to both the shLuc and shGFP constructs when no stimulation was used (1.7- to 1.9-fold). All of the TNF stimulated lines had significantly increased NFKB activity compared to all the unstimulated lines. However, there was no difference amongst the TNF stimulated samples.

It is possible that the increased baseline activity is not directly attributable to *NCSTN* haploinsufficiency. The *NCSTN* shRNA could have off-target effects, and the cells had been transduced by a virus then passaged in puromycin to generate stable knockdowns. In order to test whether this finding would repeat in another genetic model, we created an HEK293 deletion cell line using CRISPR/Cas9. We chose one single-cell dilution clone, clone 48, which had reduced *NCSTN* mRNA levels to approximately 25% of wildtype HEK293 (**Supp. Fig. 7**). We transfected the wildtype and heterozygous knockout HEK293s with the NFKB responsive luciferase, and again tested both unstimulated and TNF stimulated conditions. In this case, there is no difference in baseline NFKB activity between wildtype and knockout at baseline (**Fig. 5**; **Supp. Table 7**). As before, all of the TNF stimulated cells had increased NFKB activity compared to no stimulation. However, unlike previously the *NCSTN* deficient genome-edited line had a small but significant increase (approximately 1.75-fold; P=0.02) in NFKB activity in response to TNF stimulation.

**Fig. 4.**
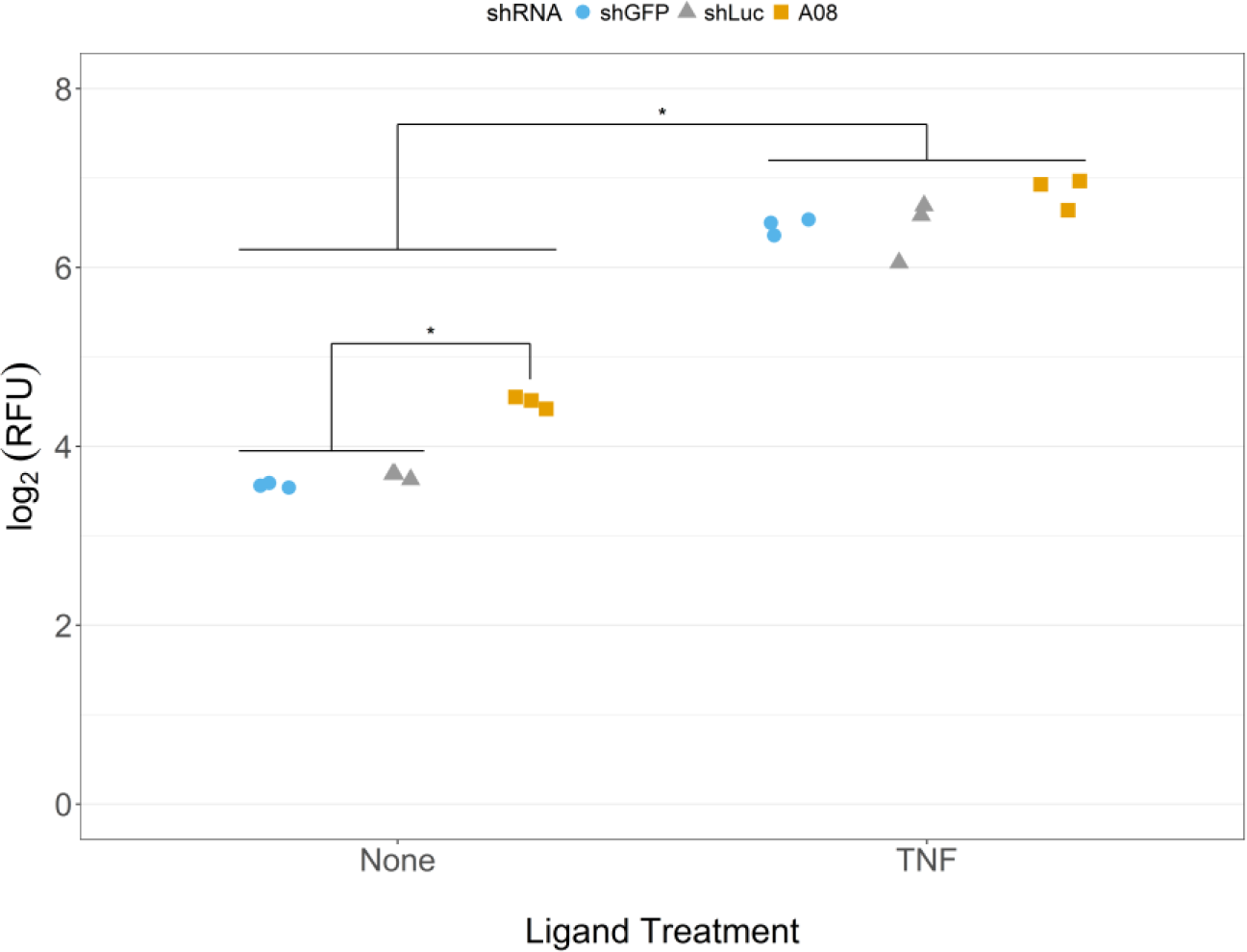
Increased baseline NFKB activity in *NCSTN* knockdowns. The x-axis shows the ligand treatment and the y-axis shows the log_2_ transformed relative fluorescence units (**RFU**). Each point represents in individual replicate. Point shape and color denote the shRNA knockdown used. Each experiment was performed in triplicate. Significant differences are shown as asterisk labeled lines.

**Fig. 5.**
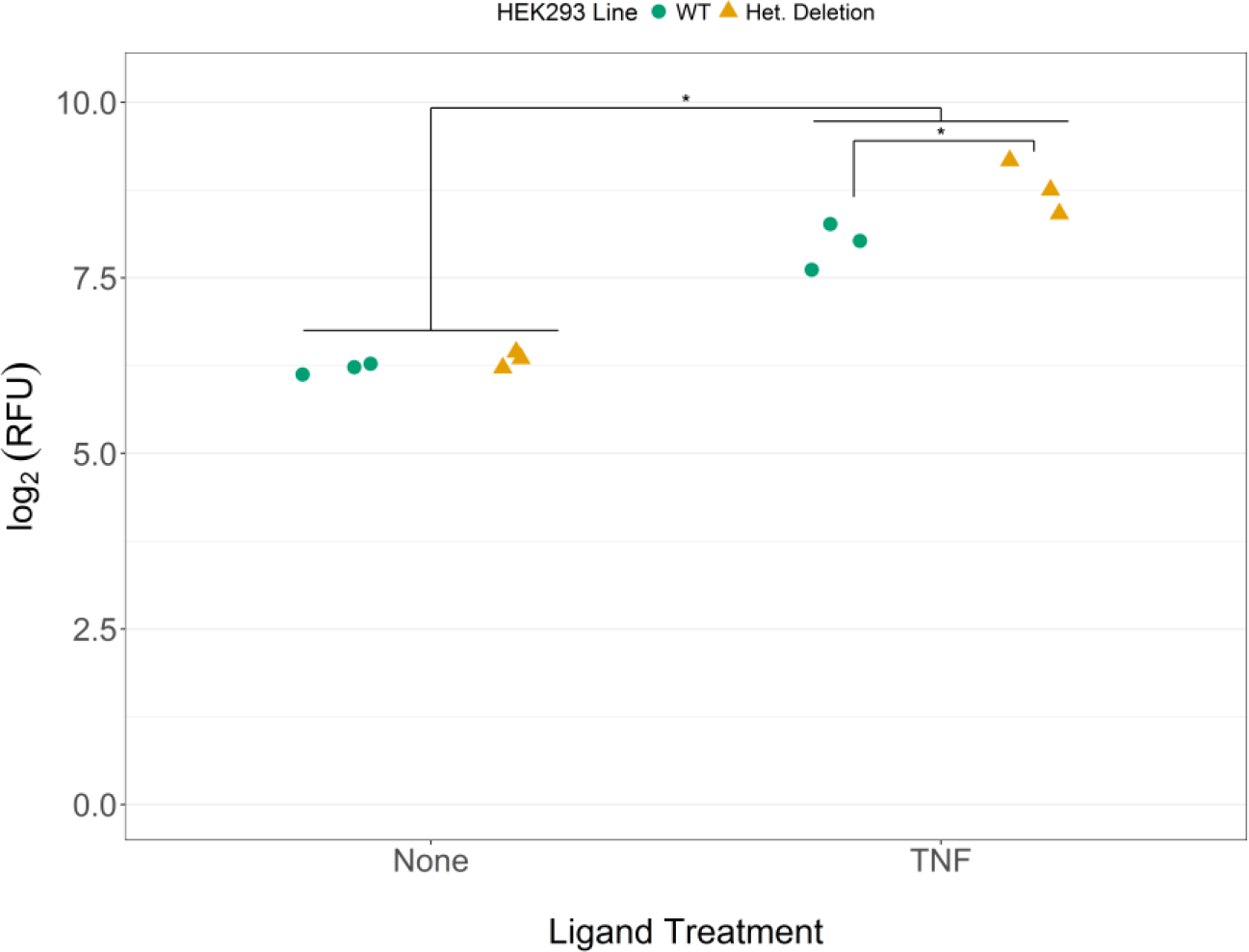
Increased NFKB activity after TNF stimulation with partial *NCSTN* genomic deletion. Results of NFKB driven luciferase activity in wildtype HEK293 (**WT**) and a partial genomic deletion line (**Het. Deletion**). The treatment is on the x-axis and the y-axis is relative fluorescence units (**RFU**). We tested triplicates of each treatment in each line, represented by the individual dots. Cell lines are differentiated by dot shape and color. Lines with asterisks indicate significant differences in activity (P < 0.05). There was no difference in baseline NFKB activity, but the deletion line has significantly increased NFKB activity after TNF stimulation

These results suggest that knockdown of *NCSTN* expression in HEK293 results in dysregulated NFKB expression. We did observe a discrepancy with regards to whether the dysregulation was observed at baseline or after TNF stimulation. This discrepancy might be a result of the intrinsic differences of the two models. It is possible that the exposure of cells to lentivirus for shRNA delivery provides a stimulus to expose the NFKB misregulation at baseline, whereas without exposure to lentivirus TNF stimulation is needed to observe the NFKB misregulation.

## E. Discussion

The natural history of HS lesion development is understudied. The basic mechanisms of pathogenesis are still controversial, with multiple competing hypotheses. The name “hidradenitis” was used because glandular inflammation was thought to be a key initiating role. However, even this may be a misnomer. It has been previously shown that an HS-like phenotype can be produced simply by applying occlusive tape, suggesting that perhaps follicular occlusion is the initiating step^27^. Histological studies have confirmed that follicular occlusion, rather than apocrinitis, is the most common feature of affected HS skin^28,29^, and that the follicles within a lesion may have weak basement membranes that are prone to rupture^30^. The rupturing of an occluded follicle alone could initiate an inflammatory response, which could be further amplified by bacterial invasion of the compromised follicle.

How would this mechanism fit with a genetic haploinsufficiency of nicastrin in familial HS? One possibility could be related to cell growth. HS lesions also epithelial hyperplasia, hyperkeratosis, and acanthosis^28,29^. In familial HS, disrupted gamma-secretase function might lead to keratinocyte overgrowth that directly contributes to occlusion of follicles. Decreased cell growth in follicular basement membranes, or otherwise weaker basement membrane structure, could promote a greater frequency of follicular rupture. Another possibility is that HS disease presentation could result from increased magnitude or duration of inflammatory response in response to an occluded follicle.

Human cell lines have obvious limitations, particularly due to chaotic rearrangements, unstable genomes, and variable ploidy. They do, however, provide a tractable system for the assay and treatment of human cells. The data we generated using these resources raise interesting questions to be pursued further in primary human cells and affected human tissue. One interesting finding is the strong increase in type-I interferon signaling in HEK001 keratinocyte knockdowns. We used a pantropic lentivirus to transduce the shRNA plasmids into the cell lines which may trigger an interferon response. However, since the effect was only seen in the keratinocytes with *NCSTN* knockdown, it must not be a general feature of lentivirus stimulation. It also was not observed in the HEK293 knockdown lines, indicating the effect must have some cell-type specificity. The interferon response also would not be in response to continued viral replication as the system we use does not produce replication-competent lentivirus. This raises the question of whether individuals with *NCSTN* haploinsufficiency have misregulated inflammatory responses. Do specific cell types in these individuals produce prolonged cytokine / chemokine signaling after initial stimulation, and could this mechanism underlie some of their disease phenotype? We have previously helped describe keratinocyte hyperinflammatory responses in individuals with familial psoriasis driven by rare variants in *CARD14*, supporting this possibility^31,32^ It will be interesting to evaluate whether local misregulated keratinocyte signaling may play a key in perpetuating familial HS lesions.

Something we did not observe in our keratinocyte lines was an equally compelling signature for NOTCH signaling. While the essential role for gamma-secretase regulation of NOTCH signaling is well-established, there is little direct evidence that *NCSTN* haploinsufficiency instigates HS disease through decreased NOTCH signaling. Furthermore, several studies have shown *in vitro* that *NCSTN* is not essential for processing all gamma-secretase substrates^33,34^. One study evaluating primary cells from individuals with *NCSTN* haploinsufficiency found no decrease in the number of gamma-secretase complexes formed in haploinsufficient cells^35^. And another study found that 3 of 4 NCSTN missense mutations associated with familial HS disease are still able to facilitate NOTCH signaling^36^. It is also worth noting that a previous array-based gene expression study comparing lesional HS skin (n=17) to non-lesional HS skin (n=13) discerned a strong signal for increased inflammation in lesional skin without a strong enrichment for NOTCH signaling^37^. In contrast, previous work using siRNA knockdown of *NCSTN* in the HACAT keratinocyte cell line did demonstrate some evidence of disrupted NOTCH signaling^38^. It also demonstrated reduced NOTCH1, NOTCH2, and NOTCH3 staining in the skin of a patient with haploinsufficiency. These findings all add to the likelihood that *NCSTN* haploinsufficiency results in a complex phenotype with tissue and time-course specific responses that may include altered NOTCH signaling, but is not solely defined by it. It is important to consider that there are many single-pass transmembrane peptides that could be cleaved by γ-secretase to produce peptide fragments with transcription activating potential. It is also possible that there are secondary functions for *NCSTN* that are independent from γ-secretase complex formation.

The finding of increased inflammatory responsiveness in keratinocyte cell lines with low *NCSTN*, increased inflammation in HS patient skin, and the effectiveness of TNF blockade in sporadic HS all suggest inflammation is key in perpetuating lesions. Additional lines of evidence support the critical role for inflammation in initiation and maintenance of HS wounds. The prevalence of HS in patients with inflammatory bowel disease is higher than general population prevalence, approaching 20% in some cohorts^11,39^. HS has also recently been shown to be more common in axial spondyloarthritis patients^40^. In one study a *TNF* promoter SNP (rs361525) was enriched in a small cohort of HS patients compared to controls^40^, and this allele also is associated with risk for psoriasis^41^. Higher levels of TNF have been observed in HS lesional skin as well^42^.

Another intriguing phenotype uncovered with our study is altered growth / cell division in cells with reduced levels of *NCSTN.* Both the HEK001 keratinocytes (decreased cell cycle markers) and the HEK293 cells (increased senescence markers) had enriched pathways suggestive of altered growth potential. Our proliferation assay results support the possibility of increased senescence or increased cell death with reduced *NCSTN.* HEK001 lines are typically more difficult to grow, and grow much more slowly, than HEK293 cells. Ultimately the HEK001 knockdowns stopped growing altogether. The HEK293 lines survived longer and allowed for some of the NFKB inducible luciferase experiments, but they also eventually stopped growing. The genome-edited HEK293 clonal line has continued to proliferate over a longer period of time. It remains to be seen whether they will eventually show a similar phenotype, or if the shRNA overexpression / selection system contributed to the effect. If this phenotype is the result of haploinsufficiency, it may suggest that individuals with low *NCSTN* might have increased rates of senescence, particularly in the pilosebaceous unit, leading to increased possibility of rupture. This result is intriguing as it is in contrast to the HACAT knockdown study, where the authors found increased proliferation in knockdown cells as assayed by CCK-8 absorbance^38^. It is worth noting that study was performed in the 12 - 72 hours post-transfection phase, whereas we checked after generating a stable knockdown line. Their results may be more specific to HACAT cell lines, or represent an early proliferation phenotype that later results in increased rates of senescence.

This study helps to highlight the potential for diverse and cell-type specific responses to decreased levels of *NCSTN.* Are there other pleiotropic effects in individuals with heterozygous loss of *NCSTN* that have familial HS? If so, some of these responses are likely to only be observed with the correct environmental stimulus and may vary from cell type to cell type. NOTCH signaling of course cannot be excluded as a driving force in pathogenesis, but inflammation may play an independent role. Indeed, there may be time-course specific progression during HS lesion development that involves NOTCH signaling initially, but eventually is driven by inflammatory stimulus. It will be important to ultimately help distinguish whether familial HS is similar to sporadic HS, sharing similar pathological mechanisms, or whether it is a severe phenocopy with unique mechanisms. Further study in patient tissue and in primary cell lines will be required to help tease apart these possibilities.

## F. Acknowledgments

Funding sources: EDOR and LC were partially supported by NIH grant P30-AR048335. DJM-H was supported by training grant T32-AR007279-36. We thank the Genome Technology Access Center in the Department of Genetics at Washington University for assistance with microarray data generation (Siteman Cancer Center grant P30CA91842 and ICTS/CTSA grant UL1TR000448). RNAi constructs were created by the Washington University RNAi core, which is supported by the Children’s Discovery Institute, The RNAi Consortium (TRC), and The Genome Institute (TGI) at Washington University. We also thank Dr. Blok for providing access to the original microarray data from their tissue study.

## H. Supplementary Data

### H.1. Supplementary Methods

#### H.1.1. Propagating standard deviation in ddCq measurements

For all of our RT-qPCR experiments, we set up at least 3 technical replicates per gene, and for each panel of tested genes included an 18S rRNA reference amplicon. We were ultimately interested in the relative expression difference of our test condition versus control condition, and calculated this metric using the “delta delta Ct” method, which we will call ddCq for quantification cycle (**Cq**) rather than threshold cycle (**Ct**). First, we calculated the mean (*x*) and standard deviation (s) for each set of technical replicates. Next, we calculated the difference between the quantification cycles (dCq) of the gene of interest and the reference gene (18S rRNA). Subtracting the means is simple, but both values have their own variability estimates (standard deviation).

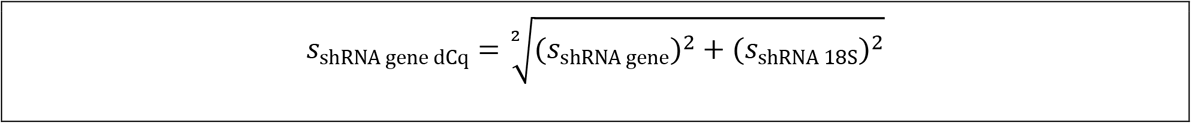

The second delta step involves solving for the difference in dCq values between the test shRNA and the control shRNA (ddCq). As before, subtracting the means is simple, and the standard deviations should be propagated similarly.

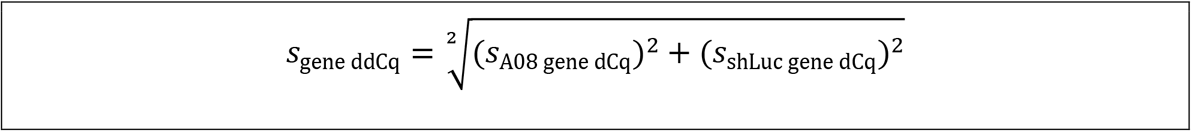

We can calculate the relative fold-change between conditions by raising 2 to the negative mean ddCq value. However, the standard deviation propagation is slightly different since we are now raising a constant to a power with error.

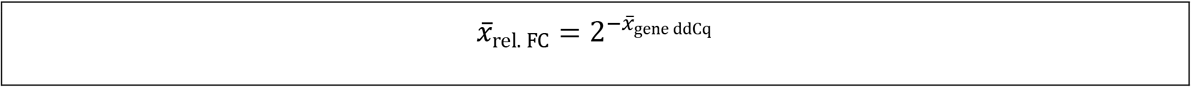

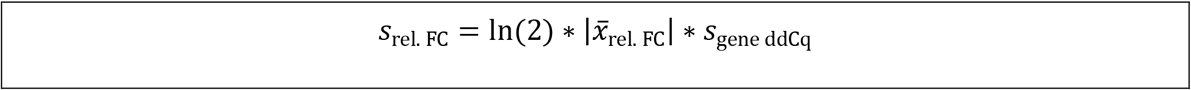

The final relative expression value therefore estimates the mean fold-change along with an error propagated measure of variability.

#### H.1.2. Propagating standard deviation for colorimetric assay fold-changes

We calculated the mean *(x)* and standard deviation (*s*) for the proliferation assay values. Calculating the fold-change between each condition therefore involves dividing the mean absorbance values.

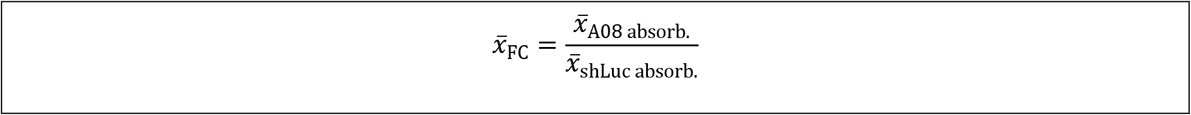

However, in this case since division is involved, we propagate the standard deviation as relative error.

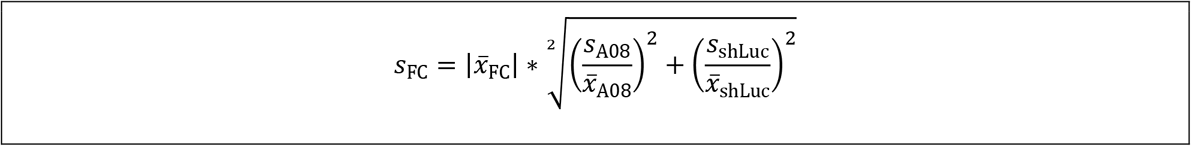

#### H.1.3. Characterizing the genomic deletion line

We screened several single-cell dilution clones after the transfection of our homology-directed repair construct, single-guide RNA plasmid, and Cas9 plasmid. Clone 48 showed a frameshifting deletion by Sanger sequencing, characterized by chaotic overlapping peaks. We amplified the target region using the *NCSTN* editing amplicon primers, and generated an amplicon sequencing library by the TruSeq method (ligate Illumina TruSeq adapters to the amplicon A-overhang). We uniquely indexed amplicons from wildtype (**WT**) HEK293 cells and the edited clone, and sequenced the amplicon as a spiked in sample on an Illumina MiSeq in 2×250 mode. Amplicon sequencing is available in the Sequence Read Archive (BioProject PRJNA268374). This generated ~168,000 reads for the wild-type cells and ~153,000 reads for the deletion clone. We cleaned the data up with cutadapt (v1.14) by removing contaminating sequencing adapter, trimming bases with qualities < 20, and requiring a minimum trimmed length of 50 bp. We aligned the cleaned data to the b37 (Broad GRCh37 + decoy contigs) with bwa mem (0.7.10-r789). Approximately 98% of all reads mapped appropriately by samtools flagstat (v0.1.19-44428cd).

We ran pindel (v0.2.5b9, 20160729) on the aligned data. It is worth noting that this version of pindel will fail to compile with errors related to use of an ambiguous absolute value function in the source code. Replacing these ambiguous assertions with an explicit alternative function (documented in the GitHub Issues tracker for pindel) will allow for successful compilation. The strongest evidence for indels included insertion of an A (GRCh37 g.1:160326443_160326444insA) in both wild-type and deletion lines. The most common deletion detected was an 8 bp deletion in the deletion clone line (GRCh37 g. 1:160326444- 160326451del). This deletion would result in a frameshift and likely non-sense mediated decay. Pindel listed many other potential deletions and insertions, but none with as much supporting evidence

We then used Bedtools coverageBed (v2.26.0-19-g6bf23c4-dirty) to find the per-base sequencing coverage within the target amplicon. This would uncover a drop in coverage in a deleted region. For both WT and the deletion line, we divided the per-base coverage by the median coverage of the whole amplicon to adjust for differences in overall depth. We then plotted the adjusted coverage across the amplicon. This plot revealed that HEK293 lines likely have an internally duplicated region or a tandem duplication of a region of our amplicon, shown by a large increase in coverage in the center of the amplicon (**Supp. Fig. 8**). However, the deletion line showed a decreased coverage over these and surrounding regions. In fact, zooming in on the coverage within the amplicon shows the likely cut site to be 4 bases internal to the single-guide RNA target, suggested successful editing. While additional work would be needed to resolve the duplicated partial copies of *NCSTN* (or partial internal duplication in HEK293s), the RT-qPCR (**Supp. Fig. 7**) combined with this data support successful editing at our target.

### H.2. Supplementary Figures

**Supp. Fig. 1.**
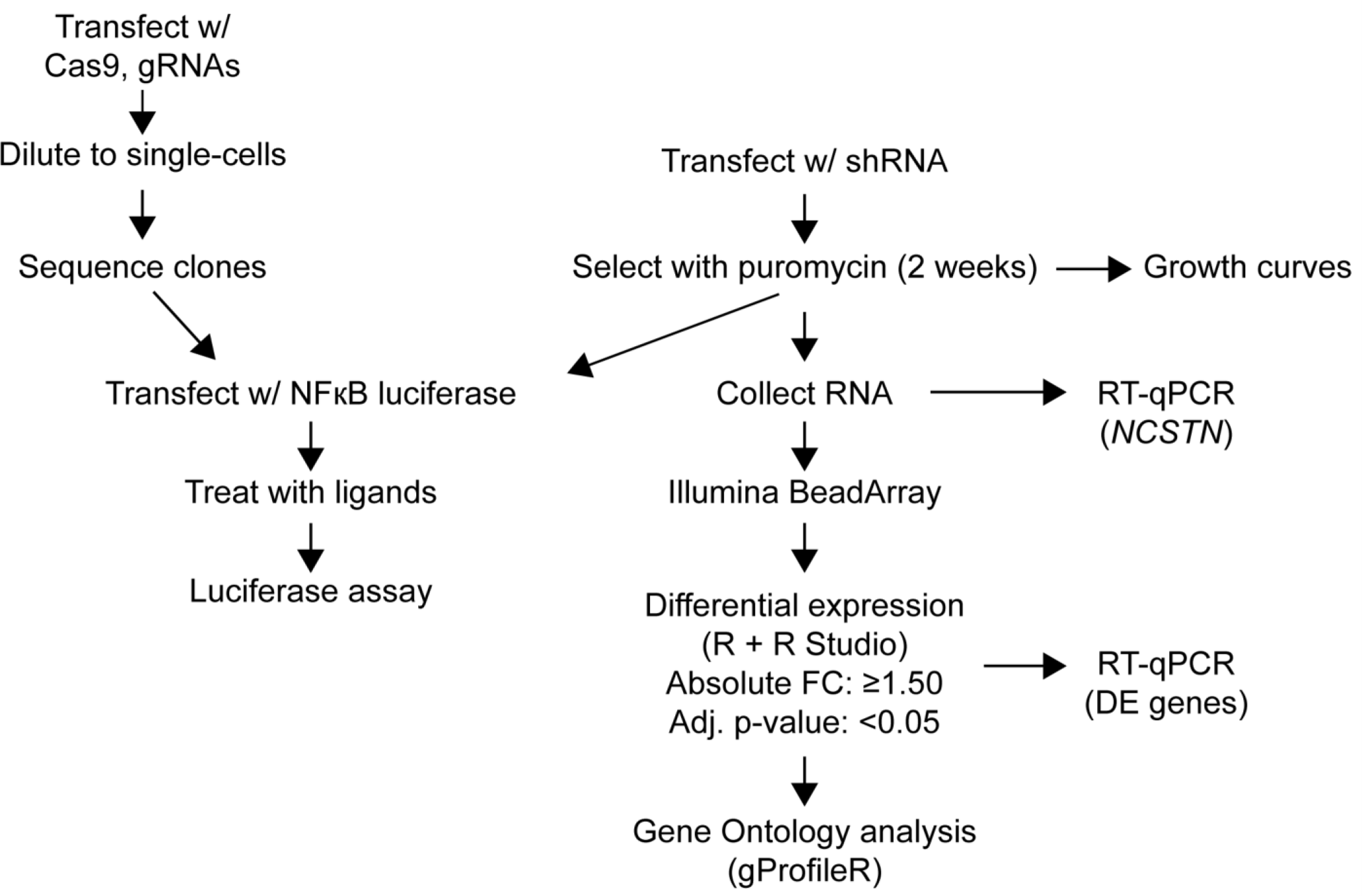
Experimental workflow. Shown above is a diagram of our approach to these experiments, including both the shRNA knockdown and genome-editing experiments.

**Supp. Fig. 2.**
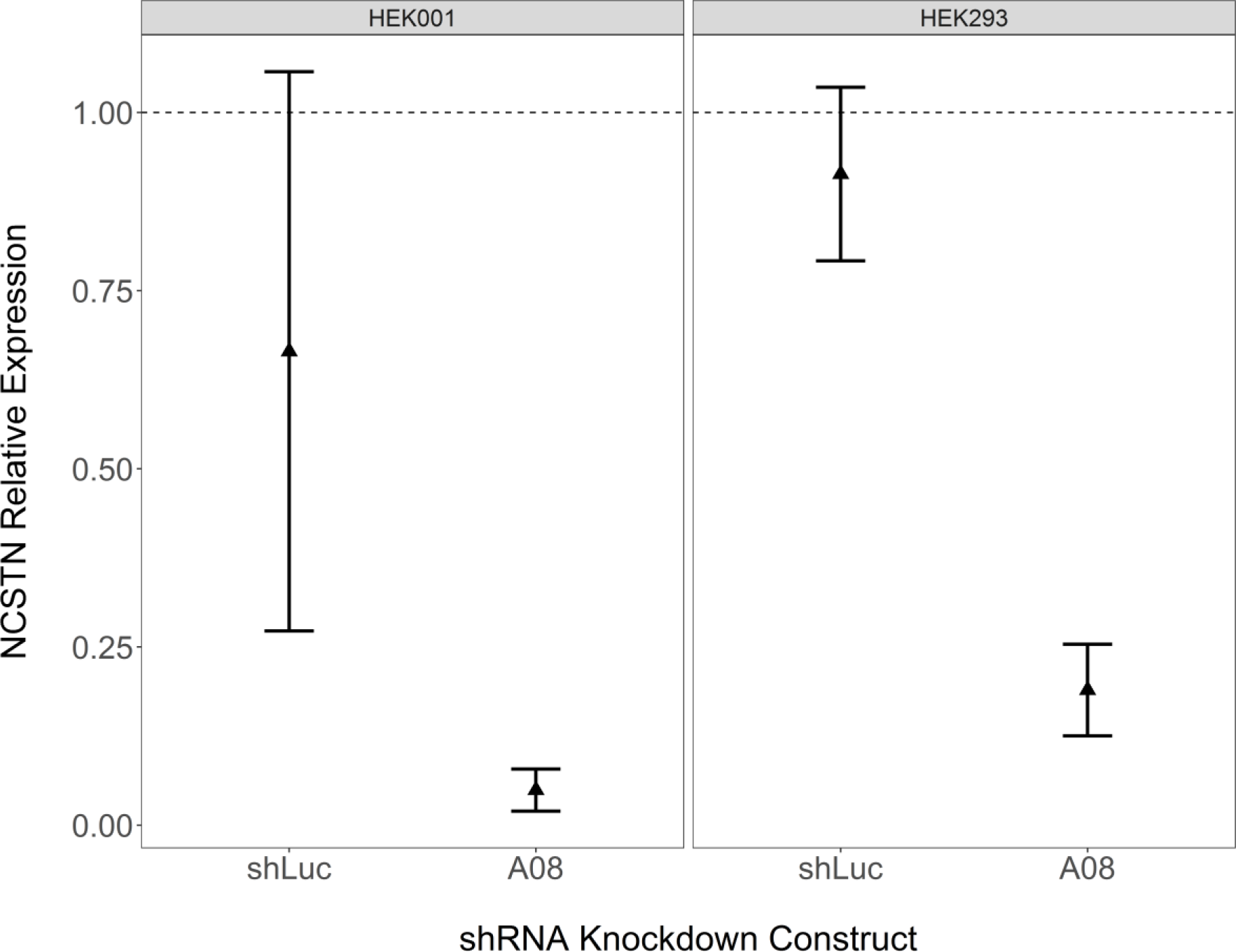
Stable knockdown confirmation. The plot shows the results for relative expression of *NCSTN* with cells knocked down with either an shRNA control (shLuc) or a *NCSTN* shRNA (A08) after 2 weeks puromycin selection. The x-axis shows the shRNA, and the y-axis shows the relative expression of *NCSTN* (amplicon 2). The points represent the mean fold-change and the error bars represent the standard deviation. Each amplicon had 3 technical replicates. The A08 shRNA had a strong knockdown in both cell lines.

**Supp. Fig. 3.**
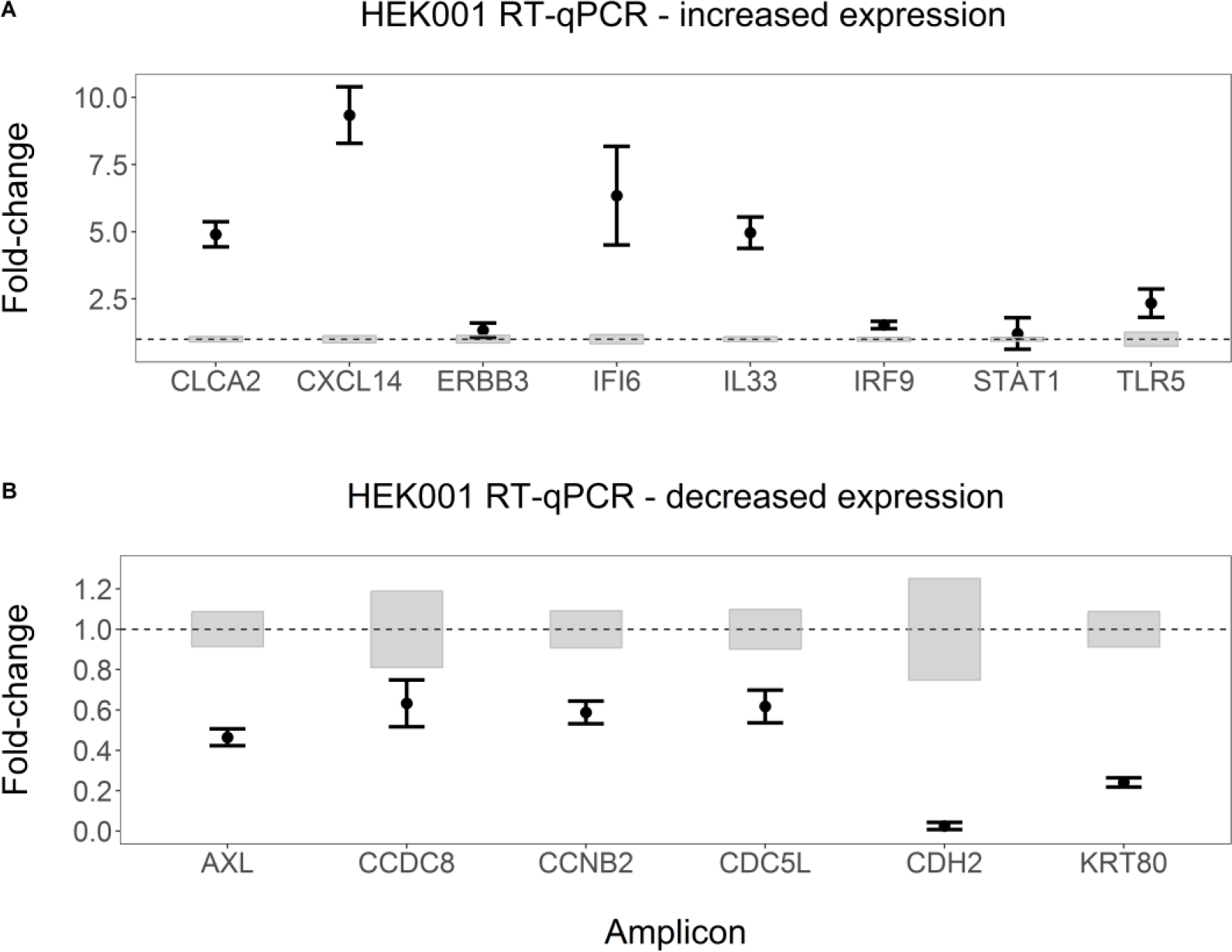
HEK001 RT-qPCR confirmation for select differentially expressed genes. Shown are the results of the confirmation studies for differentially expressed genes by microarray. Panel A shows genes with putative increased expression. Panel B shows genes with decreased expression. The x-axis shows the different tested genes. The y-axis is the estimated relative fold-change for A08 compared to shLuc. Each gene was tested in triplicate. Points indicate mean fold-change, and the error bars are the standard-deviation. For each gene, the grey box indicates the variability in that amplicon for the shLuc sample. For genes with increased expression, only *ERBB3* and *STAT1* were questionable. *STAT1* had 3 separate significant probes, suggesting that the failure to replicate may have been caused by selecting primers for the incorrect transcript. All of the decreased expression genes confirmed.

**Supp. Fig. 4.**
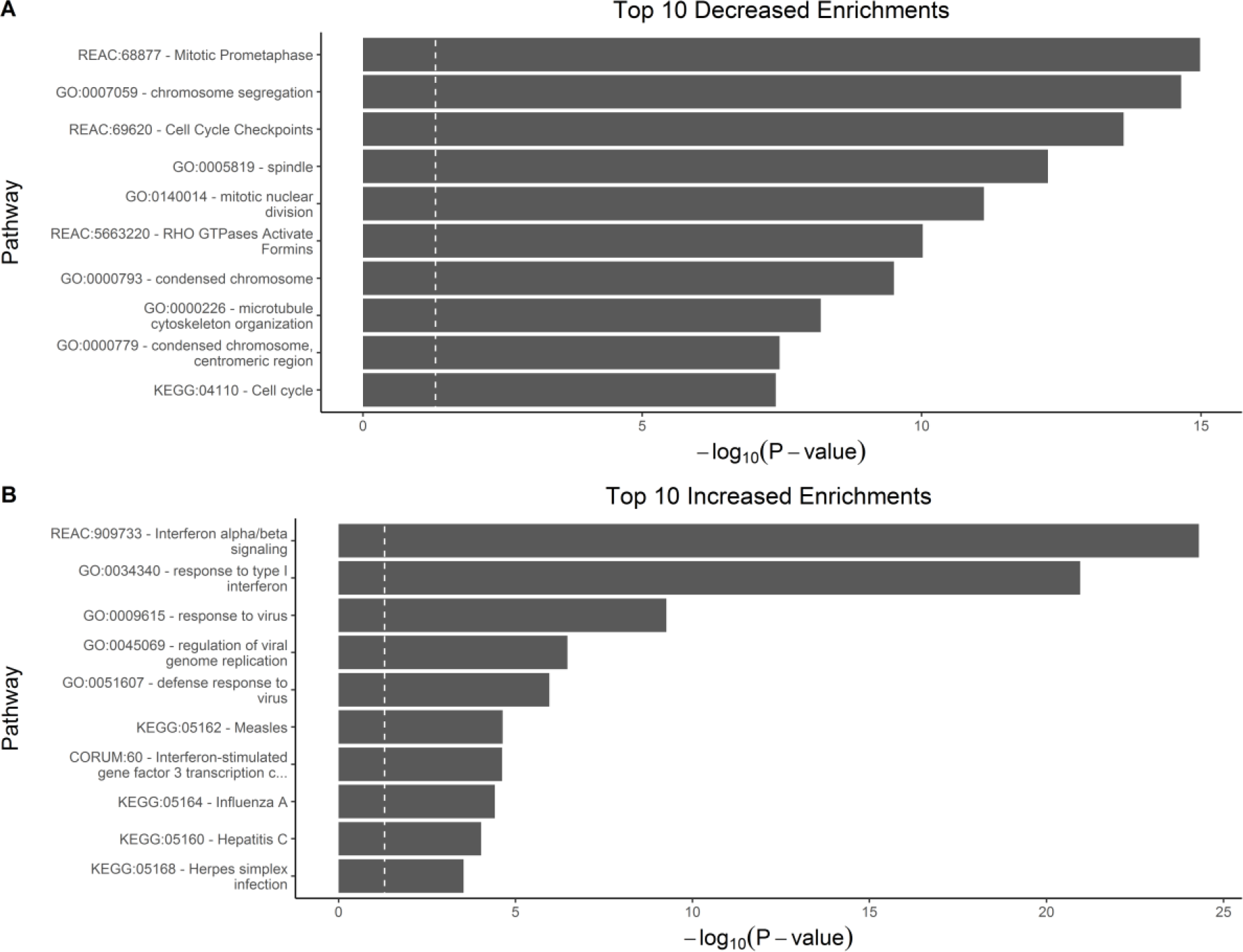
HEK001 top pathway enrichments. The x-axis shows the gProfileR significance (−log_10_ scale) for the enrichment. The white dashed line indicates the 0.05 cutoff. The y-axis shows the pathway identifier and truncated category name. Panel A shows pathways enriched amongst genes with decreased expression, and panel B are pathways enriched amongst genes with increased expression. The increased expression genes show a strong signal for type-I interferon signaling. The decreased expression genes are mostly related to cell cycle.

**Supp. Fig. 5.**
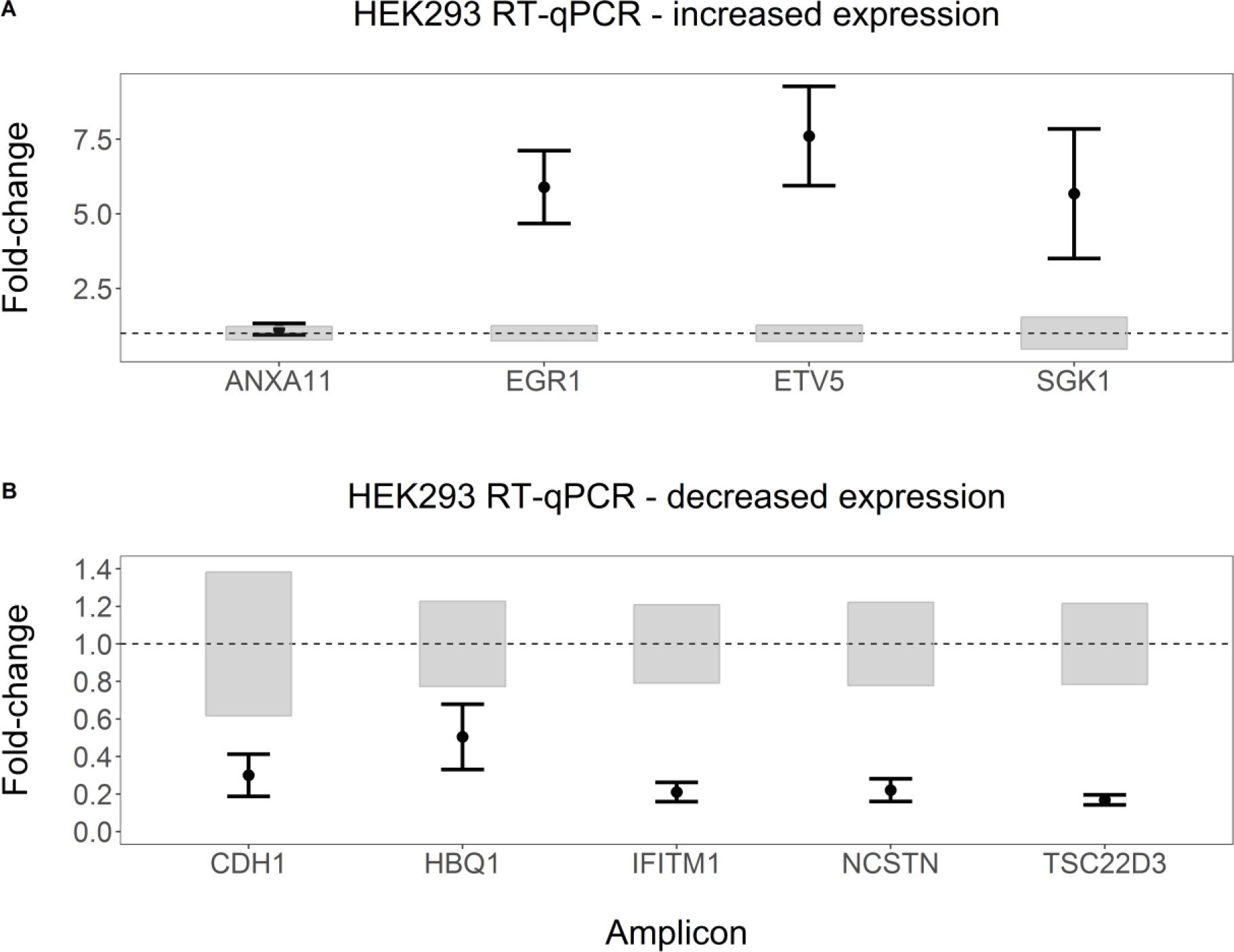
HEK293 RT-qPCR confirmation for select differentially expressed genes. Panel A shows genes with increased expression and panel B shows those with decreased expression. The x-axis shows the individual gene amplicons. Each amplicon was tested in triplicate. The y-axis is the relative fold-change of A08 knockdowns compared to shLuc knockdowns. The dots indicate mean relative expression difference, and error bars are the standard deviation. The grey boxes indicate the range of the shLuc standard deviation for that amplicon. *ANXA11* was included as a negative control and *NCSTN* as a positive control. Both behaved as expected. All of the other genes demonstrated the expected expression changes.

**Supp. Fig. 6.**
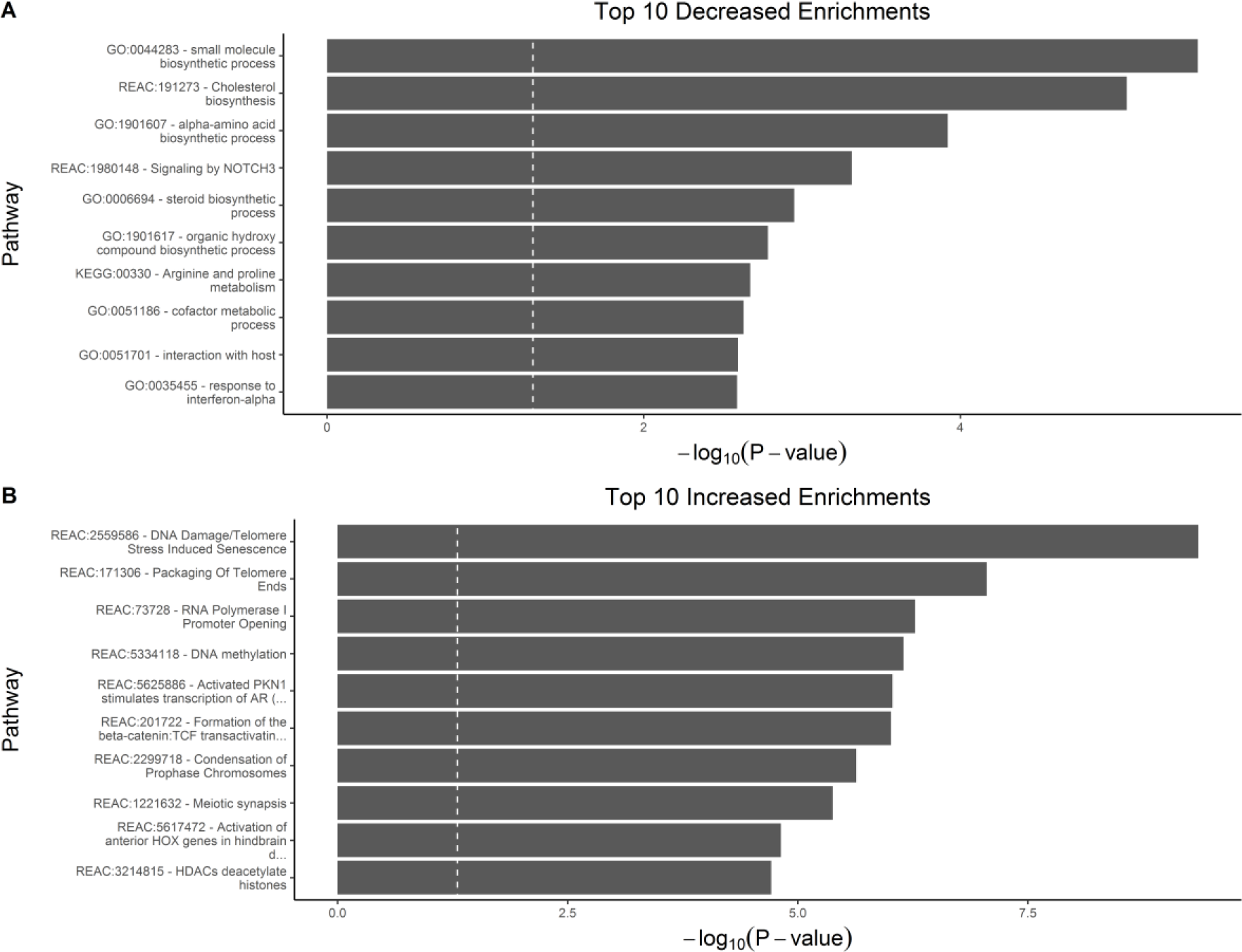
HEK293 top pathway enrichments. Shown are the top 10 pathway enrichments for genes with decreased (A) or increased (B) expression. The x-axis is the gProfileR significance (−log_10_ scale), and the y-axis is the pathway identifier along with the truncated pathway name. The decreased genes had enrichment of genes involved in cell division, such as telomere packaging and chromatin condensation. The genes with increased expression had enrichment for general metabolic processes and NOTCH3 signaling.

**Supp. Fig. 7.**
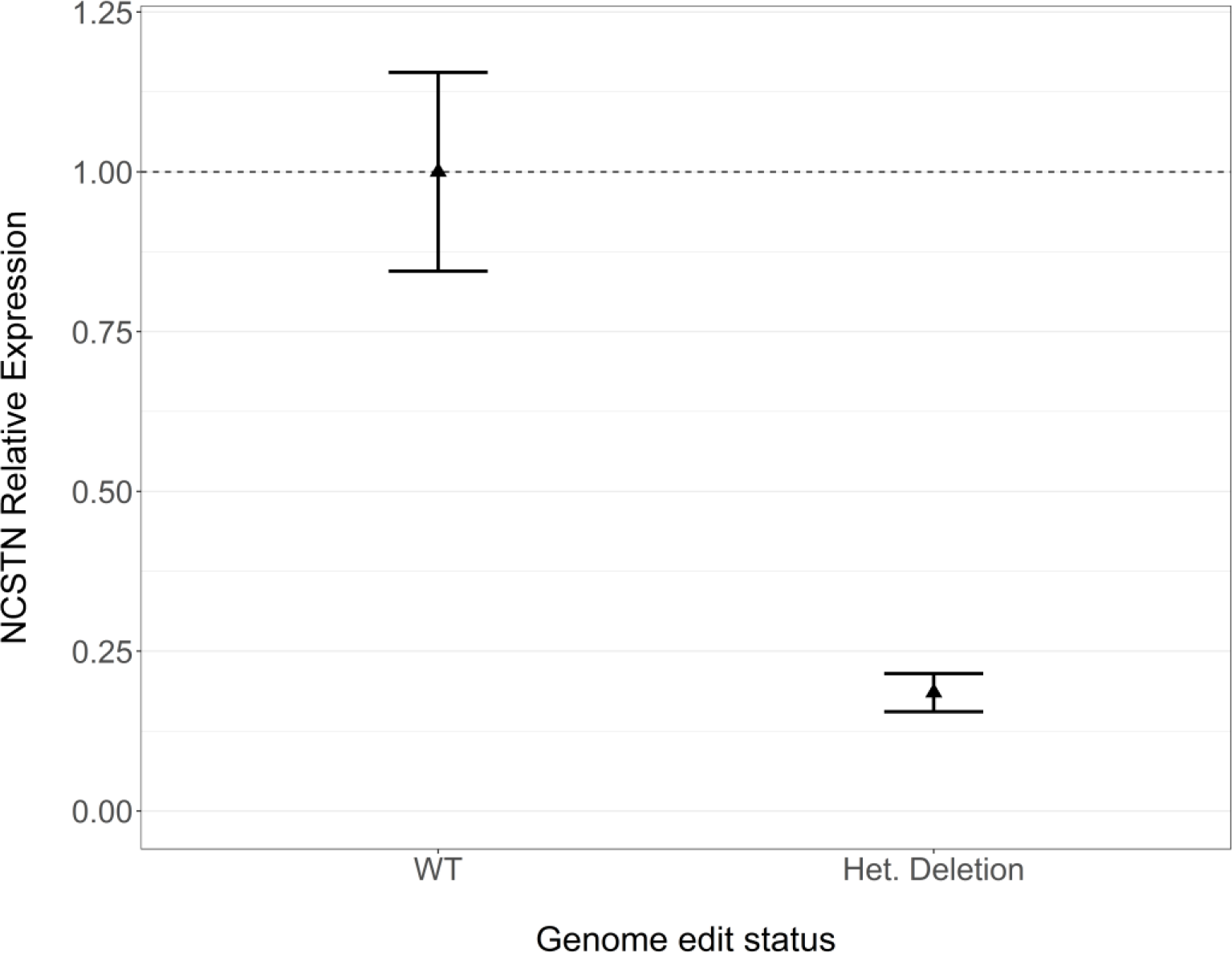
RT-qPCR validation of HEK293 engineered deletion line. Results of testing *NCSTN* expression in heterozygous deletion clone 48. The x-axis shows the cell line, where WT are unedited HEK293 cells and the Het. Deletion is the HEK293 partial *NCSTN* knockout. The y-axis is the relative expression to unedited cells based on 18S reference gene for *NCSTN* amplicon 2.The WT line is shown to give a frame of reference for the variability in WT samples. The points indicate the mean relative expression, and error bars indicate standard deviation. We measured each reference and test amplicon in triplicate for each line. The deletion clone had approximately 25% of wildtype expression, partial deletion of a multicopy locus.

**Supp. Fig. 8.**
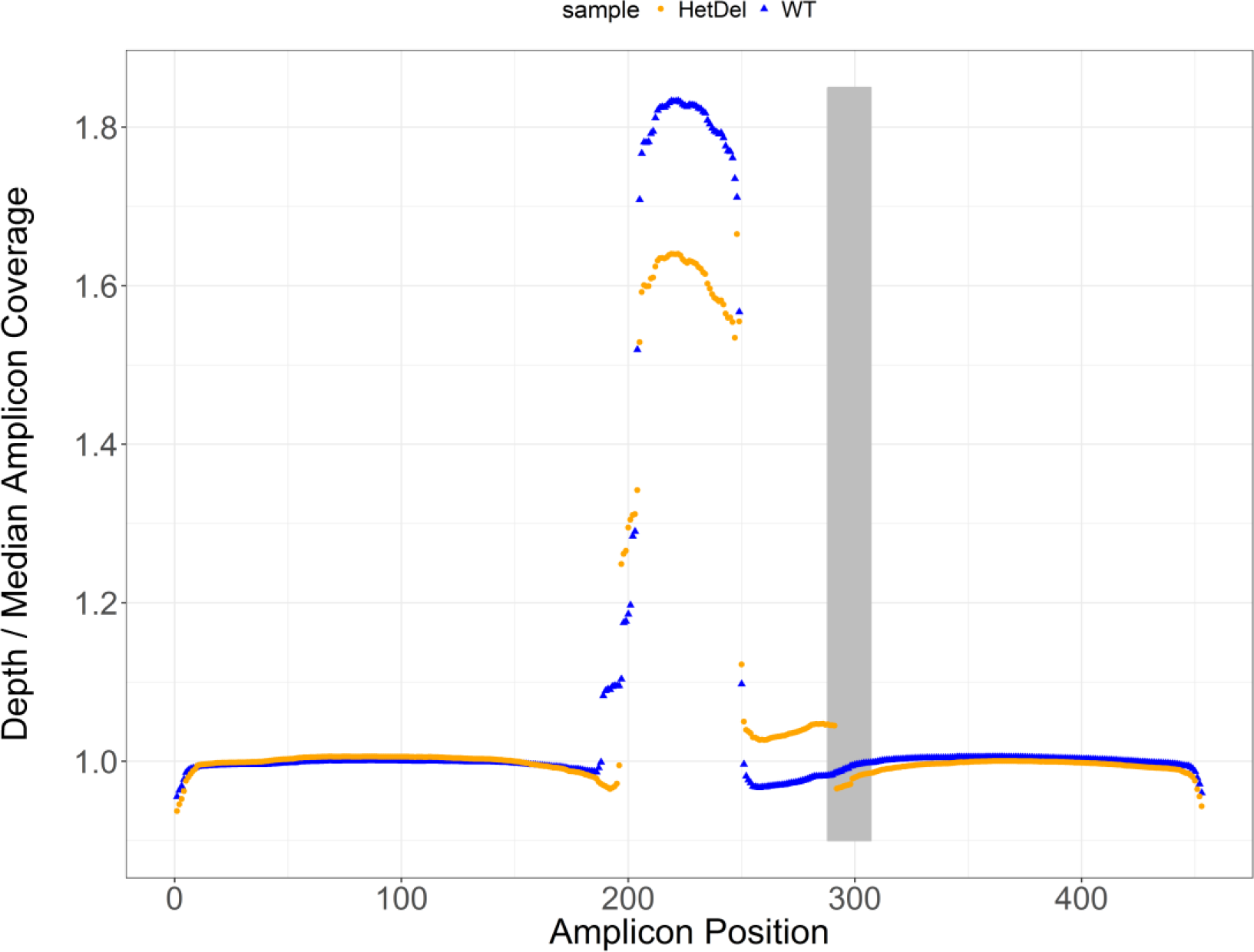
Amplicon relative coverage genome-edited line. Shown is the coverage per base of the target amplicon adjusted for total sequencing coverage. The x-axis is position within the amplicon, and the y-axis is the depth at a given site adjusted for overall aligned reads. The wildtype HEK293 line is shown in blue, and the deletion line is shown in orange. The single-guide RNA sequence is represented by the light grey rectangle. The amplicon appears to have a duplicate section from ~200-250 bp. The deletion line shows a clear drop in coverage in several regions. Additionally, the marked decrease in coverage within the sgRNA site corresponds to the expected cleavage location. This may be complicated by internal duplication during repair in the deletion line (250-300 bp), or other complex rearrangements in WT HEK293 lines.

### H.3. Supplementary Tables

**Supp. Table 1.**
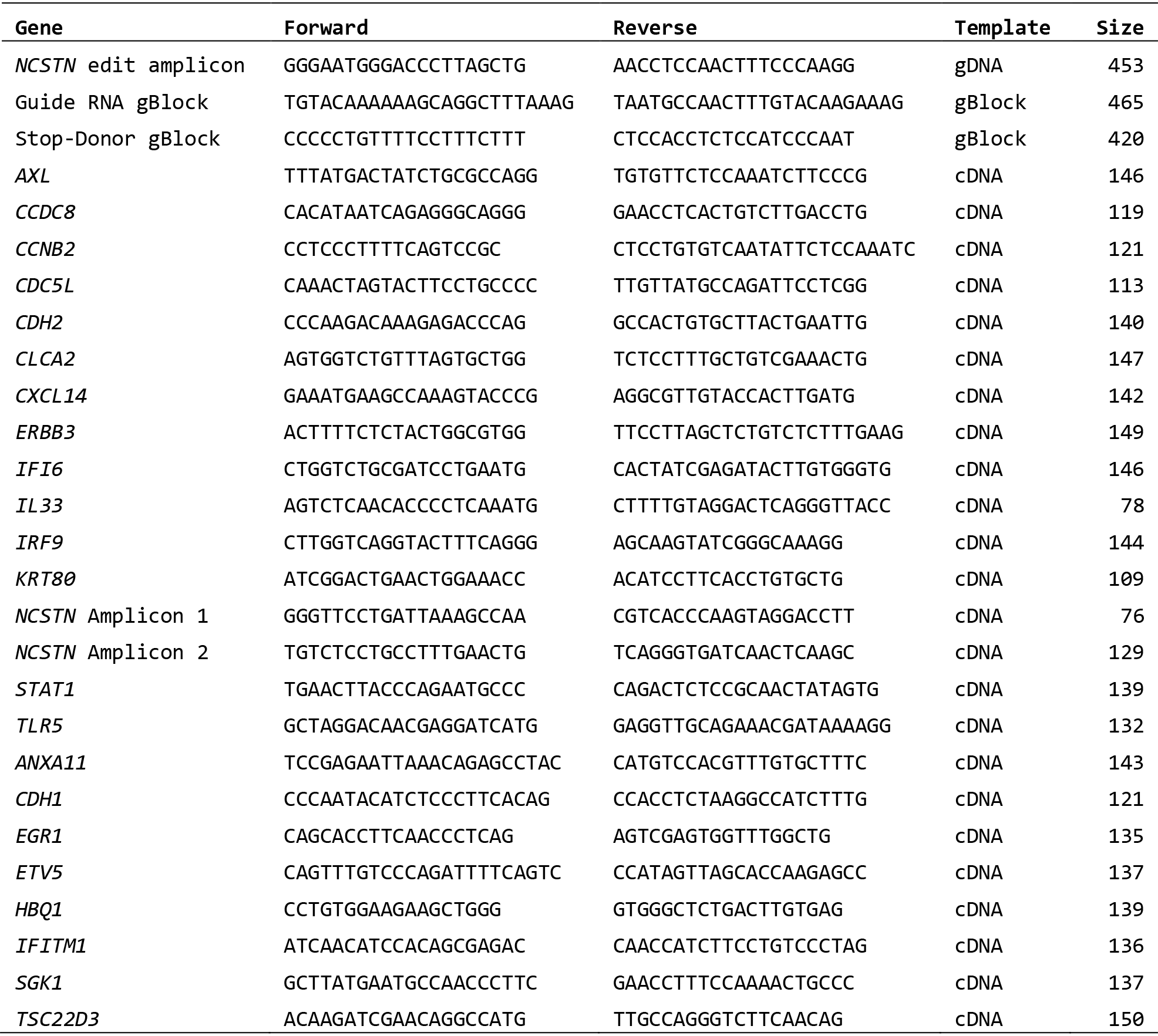
Primer sequences. Above are the gene names, primer sequences, intended template, and expected amplicon size. We determined the expected size of RT-qPCR products based on the UCSC Genome Browser In Silico PCR tool matching against UCSC genes rather than the genome.

**Supp. Table 2.**
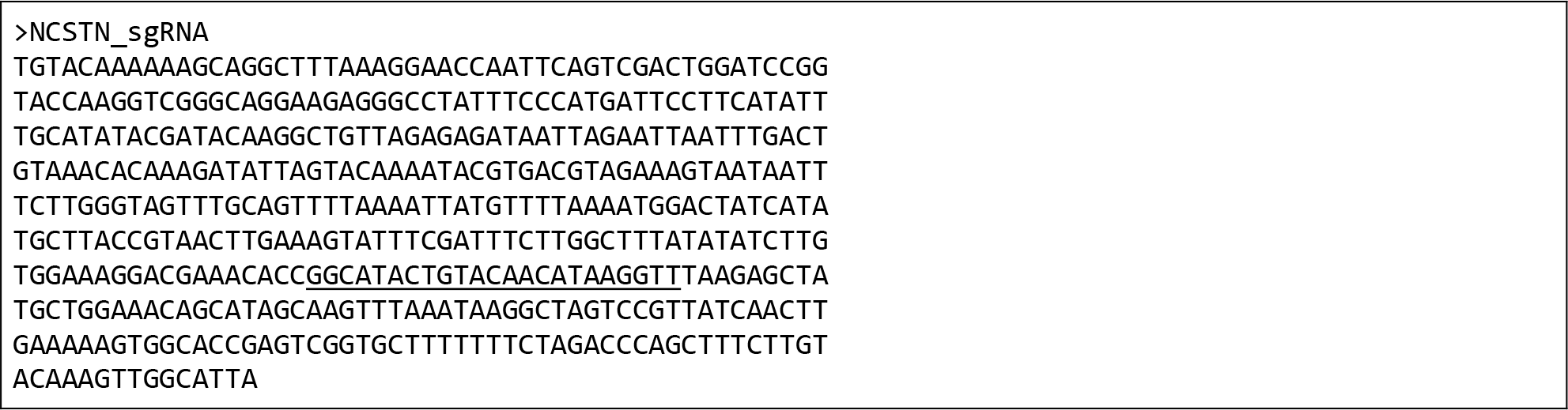
Single-guide RNA sequence. The above table contains the sequence for the single-guide RNA to target *NCSTN.* We ordered the full length sequence as an IDT gBlock and used the pCR-Blunt II-TOPO plasmid to clone it. The guide sequence is underlined.

**Supp. Table 3.**
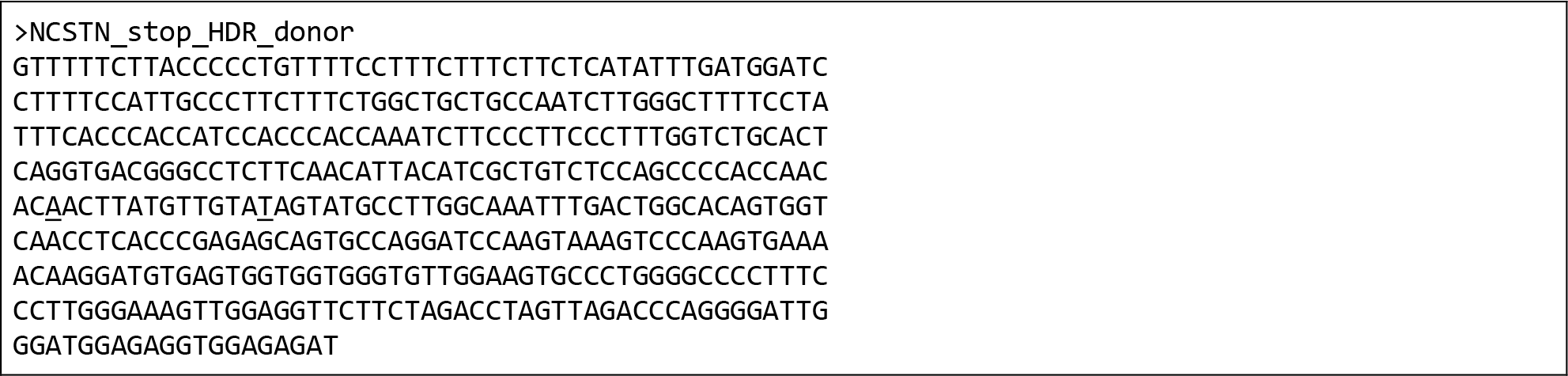
Donor sequence with stop gain. Above is the sequence for the homology-directed repair donor. We ordered this sequence as a gBIock and amplified by PCR prior to use as a donor. The donor has a mutated PAM site (first underlined sequence) to avoid being degraded by Cas9, and introduces a stop codon (second underlined base; NM_015331.2:c.1794+108C>T; p.Q568X).

**Supp. Table 4.**
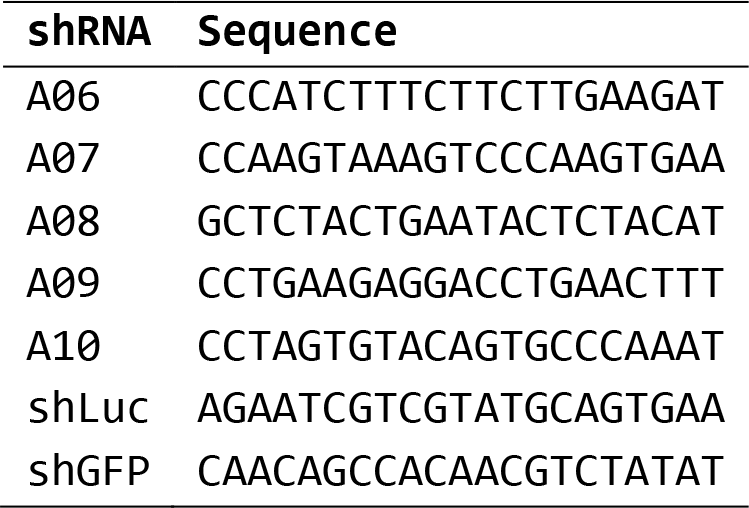
shRNA sequences. Listed are all the shRNA sequences we used in this study. All of the *NCSTN* targeting shRNAs were based on RefSeq NM_015331.1.

**Supp. Table 5.** Proliferation assay statistics. The table shows the cell lines for each of the days for colorimetric assay. We measured absorbance each day in triplicate for each cell line and knockdown treatment. We calculated the mean and standard deviation of absorbance as well. See Supplemental Methods for details of fold-change error propagation.

| Cell Line | Day | p-value | Fold change<br>(A08 / shLuc) | Fold change<br>Std. Dev. |
| --- | --- | --- | --- | --- |
| HEK001 | Day_1 | 0.5527 | 0.958 | 0.100 |
|  | Day_2 | 0.2127 | 0.917 | 0.093 |
|  | Day_3 | 0.4525 | 0.969 | 0.061 |
|  | Day_4 | **0.0428** | 0.531 | 0.116 |
|  | Day_5 | **0.0228** | 0.734 | 0.056 |
|  | Day_6 | **0.0202** | 0.652 | 0.070 |
| HEK293 | Day_1 | 0.3167 | 1.110 | 0.158 |
|  | Day_2 | 0.1162 | 1.046 | 0.041 |
|  | Day_3 | 0.9751 | 1.002 | 0.099 |
|  | Day_4 | 0.4017 | 0.957 | 0.070 |
|  | Day_5 | **0.0391** | 0.781 | 0.077 |
|  | Day_6 | **0.0434** | 0.734 | 0.085 |
|  | Day_7 | 0.0558 | 0.730 | 0.144 |

**Supp. Table 6.** NFKB luciferase activity in knockdown lines statistics. We used the Tukey Honest Simple Differences to determine estimated difference (log2 scale) and adjusted significance of the induced activity in different shRNA lines with different stimulations. The *NCSTN* knockdown line had increased baseline NFKB activity compared to both shLuc and shGFP. LHS: left-hand side. RHS: right-hand side. diff: estimated group difference with lower (lwr) and upper (upr) bounds. p adj: adjusted p-value. FC: fold-change.

| LHS |  | RHS |  | diff | lwr | upr | p adj | FC mean | FC Std. Dev. |
| --- | --- | --- | --- | --- | --- | --- | --- | --- | --- |
| shRNA | Treat | shRNA | Treat |  |  |  |  |  |  |
| A08 | None | shGFP | None | 0.93 | 0.48 | 1.39 | **1.87E-04** | 1.91 | 0.10 |
| A08 | None | shLuc | None | 0.83 | 0.37 | 1.28 | **5.75E-04** | 1.77 | 0.09 |
| A08 | TNF | A08 | None | 2.35 | 1.90 | 2.80 | **8.96E-09** | 5.10 | 0.68 |
| A08 | TNF | shGFP | None | 3.28 | 2.83 | 3.74 | **1.35E-10** | 9.72 | 1.22 |
| A08 | TNF | shGFP | TNF | 0.38 | -0.07 | 0.83 | 1.23E-01 | 1.30 | 0.18 |
| A08 | TNF | shLuc | None | 3.18 | 2.72 | 3.63 | **2.25E-10** | 9.04 | 1.15 |
| A08 | TNF | shLuc | TNF | 0.40 | -0.05 | 0.86 | 9.44E-02 | 1.32 | 0.35 |
| shGFP | TNF | A08 | None | 1.97 | 1.52 | 2.42 | **6.28E-08** | 3.92 | 0.32 |
| shGFP | TNF | shGFP | None | 2.90 | 2.45 | 3.36 | **8.07E-10** | 7.47 | 0.50 |
| shGFP | TNF | shLuc | None | 2.80 | 2.34 | 3.25 | **1.29E-09** | 6.95 | 0.48 |
| shLuc | None | shGFP | None | 0.11 | -0.35 | 0.56 | 9.65E-01 | 1.08 | 0.03 |
| shLuc | TNF | A08 | None | 1.95 | 1.49 | 2.40 | **7.14E-08** | 3.86 | 0.93 |
| shLuc | TNF | shGFP | None | 2.88 | 2.43 | 3.33 | **8.91E-10** | 7.36 | 1.75 |
| shLuc | TNF | shGFP | TNF | -0.02 | -0.48 | 0.43 | 1.00E+00 | 0.98 | 0.24 |
| shLuc | TNF | shLuc | None | 2.77 | 2.32 | 3.23 | **1.42E-09** | 6.84 | 1.63 |

**Supp. Table 7.** Increased NFKB activity with partial *NCSTN* deletion statistics. Per-comparison statistics of NFKB-driven luciferase activity in a clone genome-edited to produce lower *NCSTN.* We used Tukey Honest Simple Differences to determine estimated difference (log2 scale) and adjusted significance for our comparisons. Unlike the knockdown line, there is no difference at baseline between samples. However, activity is significantly greater in the heterozygous deletion line when both are stimulated with TNF. WT: wildtype. LHS: left-hand side. RHS: right-hand side. diff: estimated group difference with lower (lwr) and upper (upr) bounds. p adj: adjusted p-value. FC: fold-change.

| LHS |  | RHS |  |  |  |  |  | FC | FC Std. |
| --- | --- | --- | --- | --- | --- | --- | --- | --- | --- |
| Cell line | Treat | Cell line | Treat | diff | lwr | upr | p adj | mean | Dev. |
| Het. Deletion | None | WT | None | 0.13 | -0.55 | 0.81 | 9.28E-01 | 1.09 | 0.10 |
| Het. Deletion | TNF | Het. Deletion | None | 2.44 | 1.76 | 3.12 | **1.43E-05** | 5.43 | 1.49 |
| Het. Deletion | TNF | WT | None | 2.57 | 1.89 | 3.25 | **9.70E-06** | 5.93 | 1.60 |
| Het. Deletion | TNF | WT | TNF | 0.81 | 0.13 | 1.49 | **2.20E-02** | 1.75 | 0.61 |
| WT | TNF | Het. Deletion | None | 1.63 | 0.95 | 2.31 | **2.74E-04** | 3.10 | 0.75 |
| WT | TNF | WT | None | 1.76 | 1.08 | 2.44 | **1.59E-04** | 3.39 | 0.80 |

## References

1. Jemec GBE, Heidenheim M, Nielsen NH. The prevalence of hidradenitis suppurativa and its potential precursor lesions. Journal of the American Academy of Dermatology 1996; 35: 191–4.

2. Revuz JE, Canoui-Poitrine F, Wolkenstein P et al. Prevalence and factors associated with hidradenitis suppurativa: Results from two case-control studies. Journal of the American Academy of Dermatology 2008; 59: 596–601.

3. Collier F, C. SR, A. MC. Diagnosis and management of hidradenitis suppurativa. BMJ 2013; 346: f2121.

4. König A, Lehmann C, Rompel R et al. Cigarette smoking as a triggering factor of hidradenitis suppurativa. Clinical and Laboratory Investigations 1999; 198: 261–4.

5. Matusiak t, Bieniek A, Szepietowski JC. Hidradenitis suppurativa markedly decreases quality of life and professional activity. Journal of the American Academy of Dermatology 2010; 62: 706–8.e1.

6. Onderdijk AJ, van der Zee HH, Esmann S et al. Depression in patients with hidradenitis suppurativa. Journal of the European Academy of Dermatology and Venereology 2013; 27: 473–8.

7. Vinding GR, Knudsen KM, Ellervik C et al. Self-Reported Skin Morbidities and Health-Related Quality of Life: A Population-Based Nested Case-Control Study. Dermatology 2014; 228: 261–8.

8. Fitzsimmons JS, Guilbert PR. A family study of hidradenitis suppurativa. Journal of Medical Genetics 1985; 22: 367–73.

9. Pink AE, Simpson MA, Desai N et al. Mutations in the γ-secretase Genes NCSTN, PSENEN, and PSEN1 Underlie Rare Forms of Hidradenitis Suppurativa (Acne Inversa). J Invest Dermatol 2012; 132: 2459–61.

10. Schmitt JV, Bombonatto G, Martin M et al. Risk factors for hidradenitis suppurativa: a pilot study. Anais Brasileiros de Dermatologia 2012; 87: 936–8.

11. van der Zee HH, de Winter K, van der Woude CJ et al. The prevalence of hidradenitis suppurativa in 1093 patients with inflammatory bowel disease. The British journal of dermatology 2014; 171: 673–5.

12. Gao M, Wang P-G, Cui Y et al. Inversa Acne (Hidradenitis Suppurativa): A Case Report and Identification of the Locus at Chromosome 1p21.1-1q25.3. J Invest Dermatol 2006; 126: 1302–6.

13. Wang B, Yang W, Wen W et al. γ-secretase Gene Mutations in Familial Acne Inversa. Science 2010; 330: 1065.

14. Chen S, Mattei P, You J et al. γ-secretase mutation in an african american family with hidradenitis suppurativa. JAMA Dermatology 2015; 151: 668–70.

15. Li CR, Jiang MJ, Shen DB et al. Two novel mutations of the nicastrin gene in Chinese patients with acne inversa. British Journal of Dermatology 2011; 165: 415–8.

16. Liu Y, Gao M, Lv Y-m et al. Confirmation by Exome Sequencing of the Pathogenic Role of NCSTN Mutations in Acne Inversa (Hidradenitis Suppurativa). J Invest Dermatol 2011; 131: 1570–2.

17. Pink AE, Simpson MA, Brice GW et al. PSENEN and NCSTN Mutations in Familial Hidradenitis Suppurativa (Acne Inversa). J Invest Dermatol 2011; 131: 1568–70.

18. Miskinyte S, Nassif A, Merabtene F et al. Nicastrin Mutations in French Families with Hidradenitis Suppurativa. J Invest Dermatol 2012; 132: 1728–30.

19. Zhang C, Wang L, Chen L et al. Two novel mutations of the NCSTN gene in Chinese familial acne inverse. Journal of the European Academy of Dermatology and Venereology 2012; 27: 1571–4.

20. Jiao T, Dong H, Jin L et al. A novel nicastrin mutation in a large Chinese family with hidradenitis suppurativa. British Journal of Dermatology 2013; 168: 1141–3.

21. Nomura Y, Nomura T, Sakai K et al. A novel splice site mutation in NCSTN underlies a Japanese family with hidradenitis suppurativa. British Journal of Dermatology 2013; 168: 206–9.

22. Nomura Y, Nomura T, Suzuki S et al. A novel NCSTN mutation alone may be insufficient for the development of familial hidradenitis suppurativa. J Dermatol Sci 2014; 74: 180–2.

23. Kaether C, Haass C, Steiner H. Assembly, Trafficking and Function of γ-secretase. Neurodegenerative Diseases 2006; 3: 275–83.

24. Shirotani K, Edbauer D, Kostka M et al. Immature nicastrin stabilizes APH-1 independent of PEN-2 and presenilin: identification of nicastrin mutants that selectively interact with APH-1. Journal of Neurochemistry 2004; 89: 1520–7.

25. Reimand J, Arak T, Vilo J. g:Profiler—a web server for functional interpretation of gene lists (2011 update). Nucleic Acids Research 2011; 39: W307–W15.

26. Chen B, Gilbert Luke A, Cimini Beth A et al. Dynamic Imaging of Genomic Loci in Living Human Cells by an Optimized CRISPR/Cas System. Cell 2013; 155: 1479–91.

27. Shelley WB, Cahn MM. The pathogenesis of hidradenitis suppurativa in man: Experimental and histologic observations. A.M.A. Archives of Dermatology 1955; 72: 562–5.

28. Attanoose RL, Appleton MAC, Douglas-Jones AG. The pathogenesis of hidradenitis suppurativa: a closer look at apocrine and apoeccrine glands. British Journal of Dermatology 1995; 133: 254–8.

29. von Laffert M, Stadie V, Wohlrab J et al. Hidradenitis suppurativa/acne inversa: bilocated epithelial hyperplasia with very different sequelae. British Journal of Dermatology 2011; 164: 367–71.

30. Danby FW, Jemec GBE, Marsch WC et al. Preliminary findings suggest hidradenitis suppurativa may be due to defective follicular support. British Journal of Dermatology 2013; 168: 1034–9.

31. Jordan CT, Cao L, Roberson EDO et al. PSORS2 is due to mutations in CARD14. American journal of human genetics 2012; 90: 784–95.

32. Jordan CT, Cao L, Roberson EDO et al. Rare and common variants in CARD14, encoding an epidermal regulator of NF-kappaB, in psoriasis. American journal of human genetics 2012; 90: 796–808.

33. Hu C, Zeng L, Li T et al. Nicastrin is required for APP but not Notch processing, while Aph-1 is dispensable for processing of both APP and Notch. J Neurochem 2015: 10.1111/jnc.13518.

34. Hu C, Xu J, Zeng L et al. Pen-2 and Presenilin are Sufficient to Catalyze Notch Processing. J Alzheimers Dis 2017; 56: 1263–9.

35. Pink AE, Dafou D, Desai N et al. Hidradenitis suppurativa: haploinsufficiency of gamma-secretase components does not affect gamma-secretase enzyme activity in vitro. The British journal of dermatology 2016; 175: 632–5.

36. Zhang X, Sisodia SS. Acne inversa caused by missense mutations in NCSTN is not fully compatible with impairments in Notch signaling. J Invest Dermatol 2015; 135: 618–20.

37. Blok JL, Li K, Brodmerkel C et al. Gene expression profiling of skin and blood in hidradenitis suppurativa. The British journal of dermatology 2016; 174: 1392–4.

38. Xiao X, He Y, Li C et al. Nicastrin mutations in familial acne inversa impact keratinocyte proliferation and differentiation through the Notch and phosphoinositide 3-kinase/AKT signalling pathways. The British journal of dermatology 2016; 174: 522–32.

39. van der Zee HH, van der Woude CJ, Florencia EF et al. Hidradenitis suppurativa and inflammatory bowel disease: are they associated? Results of a pilot study. British Journal of Dermatology 2010; 162: 195–7.

40. Rondags A, Arends S, Wink FR et al. High prevalence of hidradenitis suppurativa symptoms in axial spondyloarthritis patients: A possible new extra-articular manifestation. Semin Arthritis Rheum 2018.

41. Savva A, Kanni T, Damoraki G et al. Impact of Toll-like receptor-4 and tumour necrosis factor gene polymorphisms in patients with hidradenitis suppurativa. British Journal of Dermatology 2013; 168: 311–7.

42. Nedoszytko B, Szczerkowska-Dobosz A, Zablotna M et al. Associations of promoter region polymorphisms in the tumour necrosis factor-alpha gene and early-onset psoriasis vulgaris in a northern Polish population. The British journal of dermatology 2007; 157: 165–7.

43. Mozeika E, Pilmane M, Nurnberg BM et al. Tumour Necrosis Factor-alpha and Matrix Metalloproteinase-2 are Expressed Strongly in Hidradenitis Suppurativa. Acta Derm Venereol 2013; 93: 301–4.

